# Isolation and Characterization of Cross-Neutralizing Coronavirus Antibodies from COVID-19+ Subjects

**DOI:** 10.1101/2021.03.23.436684

**Authors:** Madeleine F. Jennewein, Anna J. MacCamy, Nicholas R. Akins, Junli Feng, Leah J. Homad, Nicholas K. Hurlburt, Emily Seydoux, Yu-Hsin Wan, Andrew B. Stuart, Venkata Viswanadh Edara, Katharine Floyd, Abigail Vanderheiden, John R. Mascola, Nicole Doria-Rose, Lingshu Wang, Eun Sung Yang, Helen Y. Chu, Jonathan L. Torres, Gabriel Ozorowski, Andrew B. Ward, Rachael E. Whaley, Kristen W. Cohen, Marie Pancera, M. Juliana McElrath, Janet A. Englund, Andrés Finzi, Mehul S. Suthar, Andrew T. McGuire, Leonidas Stamatatos

## Abstract

SARS-CoV-2 is one of three coronaviruses that have crossed the animal-to-human barrier in the past two decades. The development of a universal human coronavirus vaccine could prevent future pandemics. We characterized 198 antibodies isolated from four COVID19+ subjects and identified 14 SARS-CoV-2 neutralizing antibodies. One targeted the NTD, one recognized an epitope in S2 and twelve bound the RBD. Three anti-RBD neutralizing antibodies cross-neutralized SARS-CoV-1 by effectively blocking binding of both the SARS-CoV-1 and SARS-CoV-2 RBDs to the ACE2 receptor. Using the K18-hACE transgenic mouse model, we demonstrate that the neutralization potency rather than the antibody epitope specificity regulates the *in vivo* protective potential of anti-SARS-CoV-2 antibodies. The anti-S2 antibody also neutralized SARS-CoV-1 and all four cross-neutralizing antibodies neutralized the B.1.351 mutant strain. Thus, our study reveals that epitopes in S2 can serve as blueprints for the design of immunogens capable of eliciting cross-neutralizing coronavirus antibodies.

## INTRODUCTION

In the past 2 decades there have been 3 zoonotic transmissions of highly pathogenic coronaviruses. SARS-CoV-1, MERS-CoV and SARS-CoV-2. The most recent one, SARS-CoV-2, has been rapidly spreading globally since late 2019/early 2020, infecting over 120 million people and killing over 2.6 million people by March 2021 (Dong et al., 2020; Patel et al., 2020). Studies conducted in mice, hamsters and non-human primates strongly suggest that neutralizing antibodies (nAbs) isolated from infected patients can protect from infection, and in the case of established infection, can reduce viremia and mitigate the development of clinical symptoms (Baum et al., 2020b; Cao et al., 2020b; Mercado et al., 2020; Rogers et al., 2020b; Schafer et al., 2021; Shi et al., 2020; Tortorici et al., 2020; Wu et al., 2020; Yu et al., 2020). Cocktails of neutralizing monoclonal antibodies have been approved by the FDA for the treatment of infection (Baum et al., 2020a; Weinreich et al., 2020). Thus, nAbs are believed to be an important component of the protective immune responses elicited by effective vaccines. Indeed, both the mRNA-based Pfizer and Moderna vaccines elicit potent serum neutralizing antibody responses against SARS-CoV-2 (Jackson et al., 2020; Walsh et al., 2020).

Monoclonal antibodies (mAbs) with neutralizing activities have been isolated from infected patients and their characterization led to the identification of vulnerable sites on the viral spike protein (S) (Cao et al., 2020a; Ju et al., 2020; Kreer et al., 2020; Liu et al., 2020; Nielsen et al., 2020; Robbiani et al., 2020; Seydoux et al., 2020; Wan et al., 2020a; Zost et al., 2020).

Many known SARS-CoV-2 nAbs bind the receptor-binding domain (RBD) and block its interaction with its cellular receptor, Angiotensin converting enzyme 2 (ACE2), thus preventing viral attachment and cell fusion (Hoffmann et al., 2020; Yan et al., 2020). However, some RBD-binding mAbs prevent infection without interfering with the RBD-ACE2 interaction (Pinto et al., 2020; Tai et al., 2020; Wang et al., 2020a). Other mAbs neutralize without binding to RBD (Chi et al., 2020; Liu et al., 2020), and their mechanisms of action are not fully understood (Gavor et al., 2020).

Plasma from SARS-CoV-1 and SARS-CoV-2 infected people contains cross-reactive binding antibodies (Ju et al., 2020; Lv et al., 2020), and a small number of monoclonal antibodies that can neutralize both viruses have been isolated from SARS-CoV-2 (Brouwer et al., 2020; Rogers et al., 2020a; Wec et al., 2020) or SARS-CoV-1-infected subjects (Tortorici et al., 2020). Overall, it appears that most of the cross-reactive antibodies do not cross-neutralize and that cross-neutralizing antibodies are infrequently generated during SARS-CoV-2 or SARS-CoV-1 infections. Antibodies capable of neutralizing SARS-CoV-1, SARS-CoV-2 and endemic human coronaviruses, such as the betacoronaviruses OC43 and HKU1 or the alphacoronaviruses 229E and NL63 have not yet been identified.

Here, we report on the isolation and full characterization of 198 S-specific mAbs from four SARS-CoV-2-infected individuals. Although a number of these mAbs recognized both SARS-CoV-2 and SARS-CoV-1, we observed minimal cross-reactivity with MERS-CoV, betacoronaviruses (OC43 and HKU1) or alphacoronaviruses (NL63 and 229E). A significant fraction of cross-reactive antibodies bound the SARS-CoV-2 S2 domain of the Spike protein. 14 mAbs neutralized SARS-CoV-2. One neutralizing mAb bound NTD, another bound the S2 subunit, one bound an unidentified site on S and the remaining 11 bound RBD. Some competed with the RBD-ACE-2 interaction while others did not. Although 7 of the SARS-CoV-2 neutralizing mAbs bound SARS-CoV-1, only 4 mAbs neutralized both viruses. Three targeted the RBD and one targeted the S2 subunit. Using the K18-hACE transgenic mouse model, therapeutic treatment with CV-30, a potent RBD-binding antibody, reduced lung viral loads and protected mice from SARS-CoV-2 infection. In contrast, a weaker anti-RBD neutralizing mAbs, CV2-75, and the anti-NTD neutralizing mAb, CV1-1, displayed minimal protective efficacies. These observations strongly suggest that neutralization potency rather than antibody epitope-specificity regulates the *in vivo* protective potential of anti-SARS-CoV-2 antibodies. Interestingly, the anti-S2 mAb, CV3-25, was the only one that was unaffected by mutations found in the recently emerged B.1.351 variant. These mAbs, especially CV3-25, can serve as blueprints for the development of immunogens to elicit protective neutralizing antibody responses against multiple coronaviruses.

## RESULTS

### Serum antibody titers and neutralizing activities against SARS-CoV-2

Peripheral blood mononuclear cells (PBMCs) and serum or plasma were collected from four SARS-CoV-2-infected adults (CV1 – previously discussed in Seydoux et al. 2020, CV2, CV3 and PCV1) at 3, 3.5, 6 and 7 weeks after the onset of symptoms, respectively **(Supplemental Table 1)**. Sera from PCV1 had the highest anti-stabilized spike (S-2P) IgG and IgM titers, while the anti-S-2P IgA titers were higher in CV1 **(Figure 1 A-C).** In contrast, to the higher anti-S-2P IgG titers in the PCV1 sera, all four sera displayed similar anti-receptor binding domain (RBD) IgG titers **(Figure 1 D-F)**. PCV1 and CV1 had higher levels of anti-RBD IgA than the other two donors and CV1 showed slightly lower anti-RBD IgM than the three other sera.

**Figure 1:**
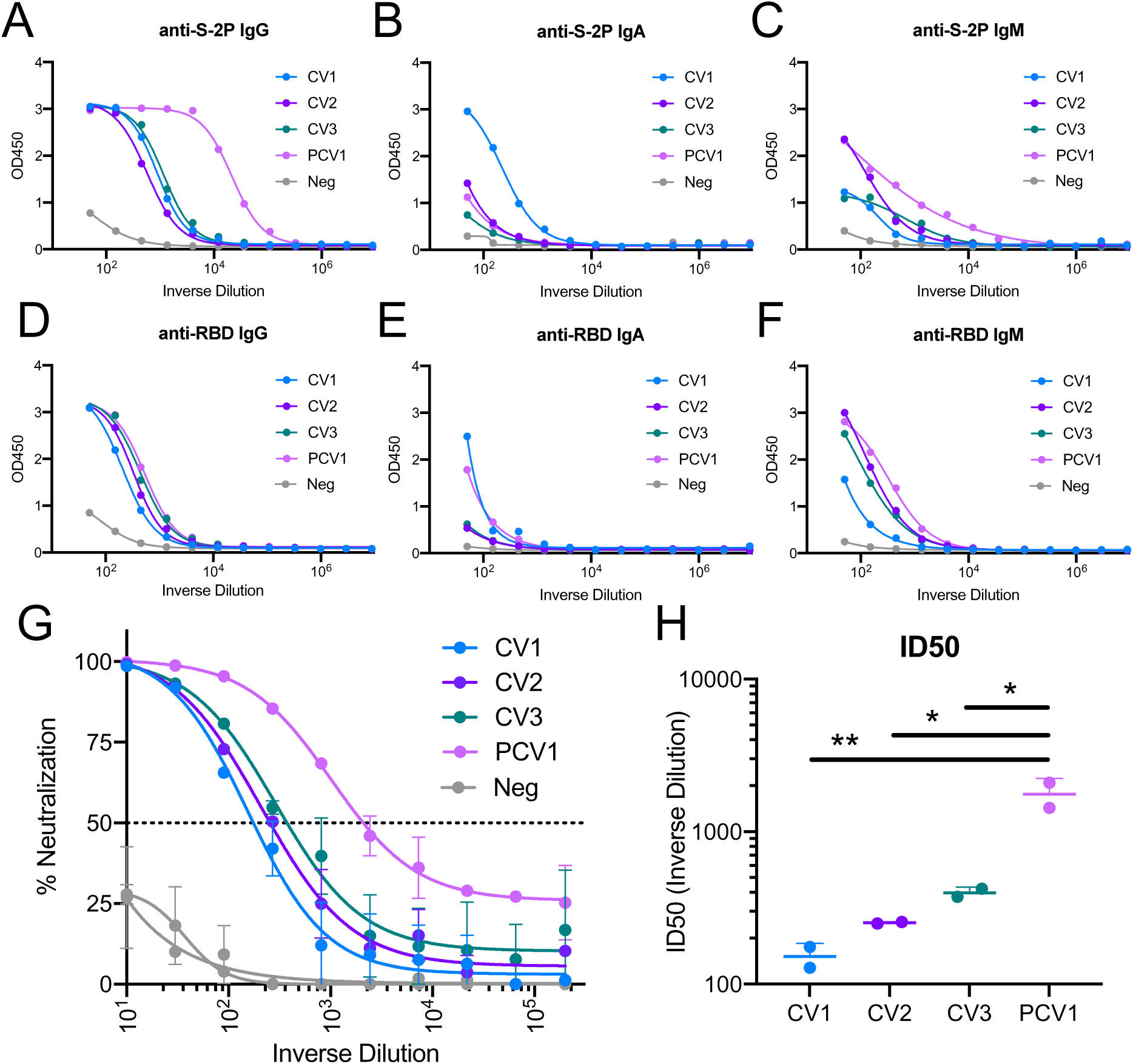
Serum antibody titers and neutralizing activities against SARS-CoV-2.

Serum from four patients infected with SARS-CoV-2 **(Supplemental Table 1)** was assessed for binding and neutralization capacity. **(A-F)** Serum antibody binding titers to S-2P and the RBD were measured by ELISA in the four participants using the indicated isotype specific secondary antibodies. CV1=Patient 1, CV2=Patient 2, CV3=Patient 3, PCV1=Patient 4. Negative sera were collected prior to the SARS-CoV-2 pandemic. **(G)** Serum from the indicated donors were evaluated for their capacity to neutralize SARS-CoV-2 pseudovirus. **(H)** ID_50_ of serum neutralization. Values are shown for two independent replicates. Statistics evaluated as one-way ANOVA with Tukey’s multiple comparison test. Significance indicated for select comparisons. *p<0.05, **p<0.01, ***p<0.001, ****p<0.0001.

While all sera neutralized SARS-CoV-2 **(Figure 1G)**, serum PCV1 was significantly more potent **(Figure 1H)**. The serum neutralizing differences do track with timepoint in infection, with the samples collected at later timepoints show greater potency, potentially indicating maturation of the humoral response. Thus, though all four patients had similar anti-RBD binding antibody titers, PCV1 developed higher anti-S-2P binding antibody titers and higher neutralizing titers than the other three patients examined here.

### Specific VH and VL genes give rise to anti-S antibodies during SARS-CoV-2 infection

Monoclonal antibodies (mAbs) have been isolated and characterized previously by us and others (Cao et al., 2020a; Ju et al., 2020; Kreer et al., 2020; Nielsen et al., 2020; Robbiani et al., 2020; Seydoux et al., 2020). We isolated individual S-2P+ and RBD+ IgG+ B cells **(Supplemental Table 1)** from all four subjects. The percentage of S-2P+ cells in the four patients ranged from 0.23%-1.84% of IgG+ B cells and out of which 5-12.7% targeted the RBD. In agreement with the above-discussed serum antibody observations, the frequency of S-2P+ IgG+ B cells in PCV1 was 3-8-fold higher than those in the other patients while no major differences were observed in the frequencies of RBD+ IgG+ B cells among the four patients. As expected, the frequency of S-2P+ cells in a healthy (pre-pandemic) control individual was lower than those found in the four patients (0.104% and 0.128%), as were the frequency of RBD+ IgG+ B cells (first sort: 0.015% and second sort: 0.019%). A total of 341 HC, 353 κLCs and 303 λLCs were successfully sequenced from the four SARS-CoV-2-positive donors **(Supplemental Table 1, Supplemental Figure 1)**, from which 228 paired HC/LCs were generated, and 198 antibodies were successfully produced and characterized. 59 mAbs were generated from the healthy individual. As discussed above we performed an initial characterization of the 45 mAbs from CV1 (Seydoux et al., 2020), here we performed a more in-depth characterization of these mAbs.

In agreement with previous reports, the antibodies isolated from the patients utilized diverse V regions (Cao et al., 2020a; Nielsen et al., 2020; Robbiani et al., 2020; Seydoux et al., 2020) **(Figure 2A-C, Supplemental Figure 1)**. Similarly, the S-specific mAbs isolated from the healthy donor originate from diverse V regions. To determine whether anti-S-2P+ B cells that express certain VH and VL genes preferentially expand during infection, we compared the relative frequencies of each VH and VL sequence to those present in healthy individuals. For this, we performed a 10x-based sequence analysis of total circulating B cells (i.e., not S-2P specific) from 5 SARS-CoV-2-unexposed adults (**Figure 2D-F, Supplemental Figure 2)**. Significantly higher frequencies of S-2P+ IGHV3-30 and IGHV1-18 antibody sequences were observed in the patients as compared to the relative frequencies of these two genes present in healthy adults (**Figure 2D**). Interestingly, lower frequencies of S-2P+ IGHV3-33 usage was observed in the patients than in healthy donors. Differences were also observed in kappa (**Figure 2E**) and lambda (**Figure 2F**) gene usage between patients and healthy donors. Specifically, IGKV3-15, IGKV1-33/1D-33 and IGKV1-17 were significantly elevated in patients as compared to healthy donors while the expression of IGKV1-39/1D-39 was reduced. IGLV1-51 was significantly elevated in the patients as compared to healthy donors, as was IGLV2-23, though this appears to be driven by a greatly elevated usage in patient CV3.

**Figure 2:**
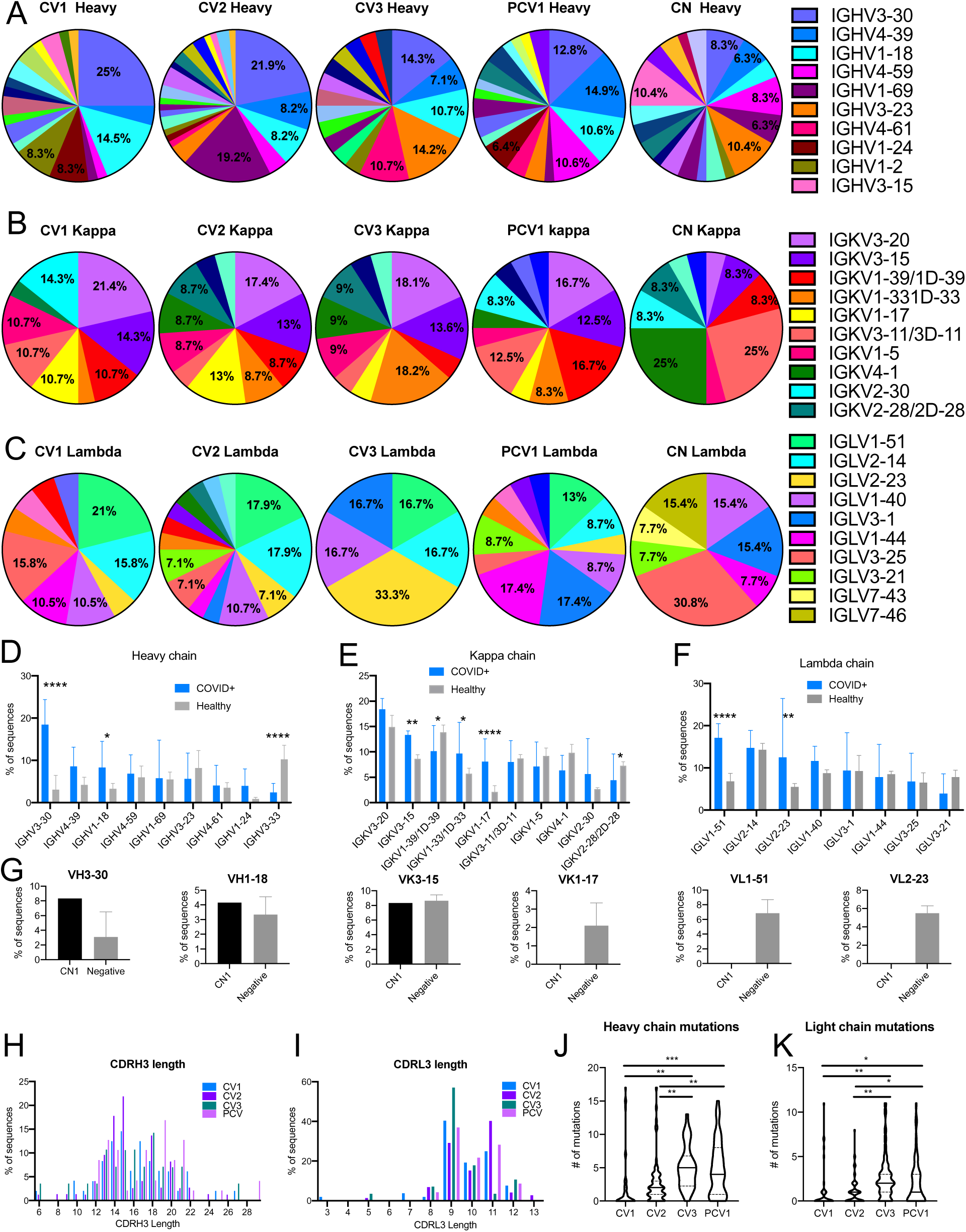
Specific VH and VL genes give rise to anti-S antibodies during SARS-CoV-2 infection. Sequences for the 198 mAbs elicited from the SARS-CoV-2 infected patients were compared for VH and VL gene usage **(A-C)**. The V gene usage was assigned for all paired heavy, **(A)**, kappa, **(B)**, and lambda, **(C)** chains recovered from S-2P-specific B cells. Percentages are shown on graph for V chains that make up more than 5% of the total for each sort. Full sequencing data in **Supplemental Figure 1**. **(D-F)** The frequency of select heavy **(D)**, Kappa **(E)**, and Lambda **(F)** chain V gene usage for the four COVID+, S-2P+ sorted participants is compared to 5 SARS-CoV-2 unexposed, ‘healthy’ adult participants determined using unbiased 10X sequencing of total B cells. Full sequencing in **Supplemental Figure 2A**. **(G)** Comparison of VH3-30, VH1-18, VK3-15, CK1-17, VL1-51, and VL2-23, frequencies in S-2P+ sorted unexposed cells (CN) and B cells from 5 unexposed donors determined by unbiased sequencing (Negative). Full sequencing in **Supplemental Figure 2B**. **(H, I)** The CDR3 length distribution for the heavy **(H)** and light chains **(I)** shown as percentage of antibodies from each donor. **(J, K)** The number of amino acid mutations in heavy **(J)** and light chains **(K)** of paired mAb sequences. Median indicated as a solid line with quartiles indicated in dashed lines. Significant differences were determined using one-way ANOVA with Tukey’s multiple comparison test *p<0.05, **p<0.01, ***p<0.001, ****p<0.0001.

The above observations suggest that naïve B cell clones expressing the above IGHV, IGKV or IGLV genes preferentially recognize the viral S protein at the initial stages of infection. To address this point IgD+ IgM+ S-2P+ and RBD+ B cells were isolated from a healthy donor, CN1 (following two independent B cell sorting experiments from this donor) **(Supplemental Table 1)**, their V genes sequenced, and their relative frequencies were again compared to those found in total B cells from healthy donors **(Figure 2 G)**. Although IGHV3-30 was present in higher frequency in B cells sorted with S-2P from CN than in the total B cell population, the difference was not as large as in the infected patients. Similarly, no differences were observed for the other IGHV, IGHV1-18 and no instances of the IGKV or IGLV genes that were predominant in the anti-S response after infection appeared in CN. Thus, it appears that the anti-spike B cell response that predominates at 3-7 weeks post infection is dissimilar from the naïve B cells that preferentially bind to S-2P.

The length distribution for the CDRH3 and CDRL3 of antibodies isolated after infection were comparable to those present in the pre-infection, healthy B cell repertoires (**Figure 2H, I**). Interestingly, the IGHVs and IGLVs sequences derived from samples collected at 6 (CV3) and 7 (PCV1) weeks after infection had significantly more amino acid mutations than those derived from samples collected at 3 (CV1) or 3.5 (CV2) weeks after symptom-development (**Figure 2J, K**). These observations are suggestive of a continuous B cell evolution during SARS-CoV-2 infection as others recently reported (Gaebler et al., 2021).

### Epitope-specificities and cross-reactivities of SARS-CoV-2 antibodies

The binding specificities of the 198 monoclonal antibodies (mAbs) to S-subdomains were determined using recombinant proteins including, S1, RBD, N-terminal domain (NTD) and S2 ectodomain (ECD) subunits **(Figure 3A, Supplemental Figure 3 A, B)**. Only a small percentage of mAbs bound RBD, irrespective of the time of B cell isolation following the development of symptoms. However, the relative proportion of anti-S2 antibodies was higher in samples collected at 3 and 3.5 weeks (51% in CV1 and 70% in CV2, respectively) than in samples collected 6- and 7-weeks post symptom onset (35% in CV3 and 27% in PCV1, respectively). PCV1 had a high proportion (32%) of antibodies whose epitopes could not be mapped to S1 or S2, while such antibodies were rarer in the other three patients examined here (15% in CV1, 7% in CV2 and 0% in CV3).

**Figure 3:**
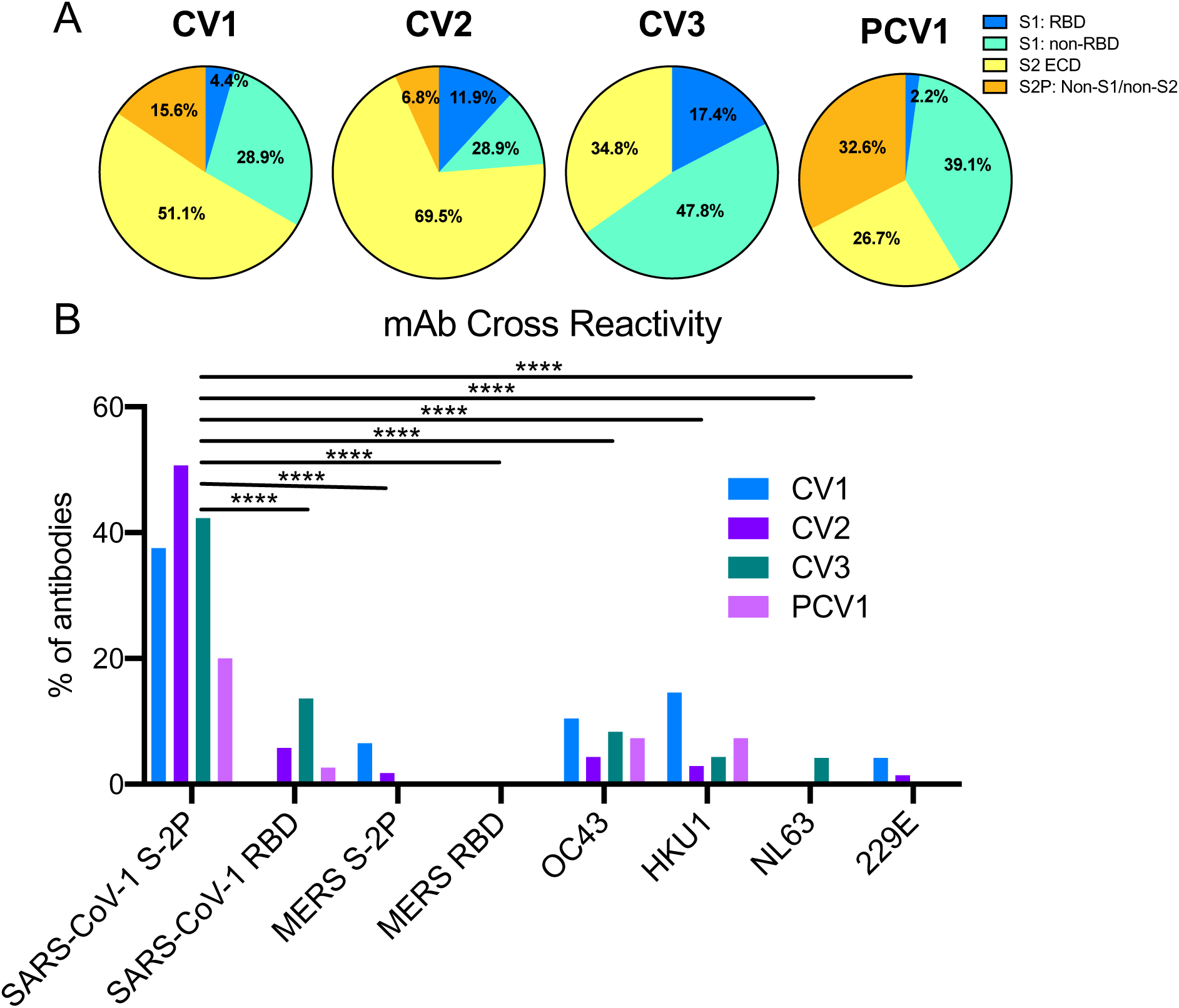
Epitope-specificities and cross-reactivity of SARS-CoV-2 antibodies. The percentage of mAbs from each donor specific for the SARS-CoV-2 spike subdomains and their cross-reactivity was determined by BLI. **(A)** mAbs were grouped into the antibodies that bound RBD in the S1 subunit (S1: RBD, blue), mAbs that bound S1 outside of RBD (S1: non-RBD, teal), mAbs that bound the S2 ECD (S2 ECD, yellow) or those that bound S2P but did not bind either S1 or S2 (S2P: Non-S1/Non-S2. **(B)** The percentage of mAbs that bind to SARS-CoV-1, MERS and the four common human coronavirus was also measured by BLI. Significant differences were determined using two-way ANOVA with Tukey’s multiple comparison test, *p<0.05, **p<0.01, ***p<0.001, ****p<0.0001. Additional BLI data and comparison to number of amino acid mutations in **Supplemental Figure 3**.

We also determined the abilities of these antibodies to recognize SARS-CoV-1, MERS, the two endemic human beta coronaviruses, OC43 and HKU1, and the two endemic human alpha coronaviruses, NL63 and 229E **(Figure 3B)**. 81 mAbs (41%) displayed SARS-CoV-1 reactivity (to varying degrees), approximately half of which recognized the SARS-CoV-1 RBD. In contrast, only 4 mAbs (2.3%) displayed cross reactivity towards MERS (and none to the MERS RBD), 13 bound OC43 (7.1%), 12 bound HKU1 (6.6%), 2 bound NL63 (1.1%) and only 1 bound 229E (0.56%). There was no association between the number of amino acid mutations in the antibody V regions and cross-reactivity with divergent HCoVs **(Supplemental Figure 3C-I)**.

### SARS-CoV-1 and SARS-COV-2 Cross-neutralizing properties of mAbs

Only 14 mAbs (7%) neutralized SARS-CoV-2 **(Figure 4A)**, with IC_50_s ranging from 0.007 μg/ml to 15.1 μg/ml (although, as we discuss below, we were unable to assign an IC50 to CV2-74) **(Figure 4B, Supplemental Table 2, Supplemental Figure 4)**. 11 of 14 neutralizing mAbs bound RBD, in agreement with our (Seydoux et al., 2020) and other reports that RBD is the major target of anti-SARS-CoV-2 nAbs (Barnes et al., 2020; Cao et al., 2020a; Ju et al., 2020; Liu et al., 2020; Rogers et al., 2020a). Three of the nAbs, CV1-1 (from patient CV1), CV2-74 (from patient CV2) and CV3-25 (from patient CV3) bound epitopes outside the RBD. CV1-1 binds the S1 NTD, CV3-25 binds the S2 subunit, CV2-74 bound neither the recombinant S1 or S2 proteins used here, and we were unable to define its specificity **(Figure 5B and Supplemental Figure 4A, B)**.

**Figure 4:**
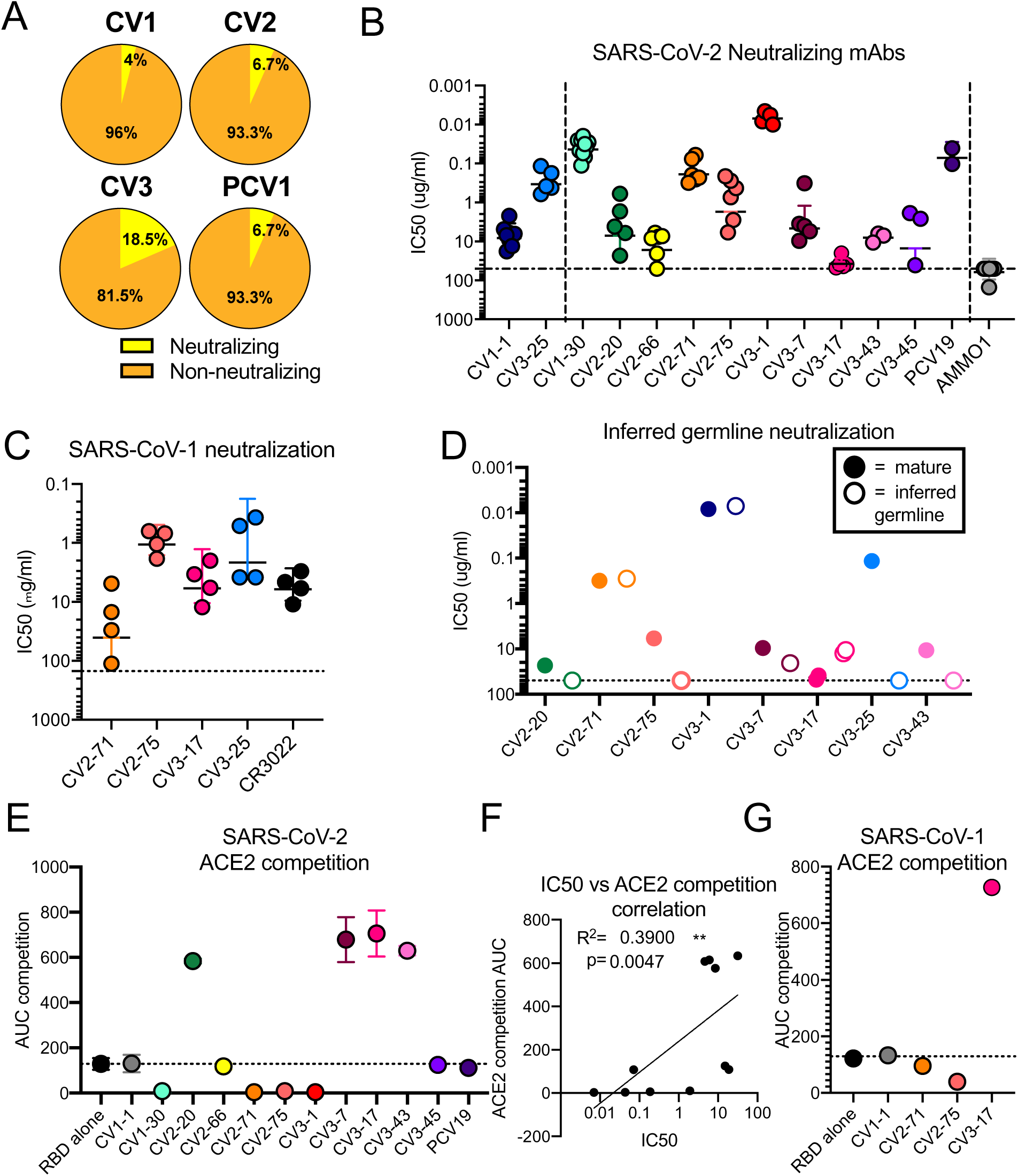
SARS-CoV-1 and SARS-COV-2 Cross-neutralizing properties of mAbs. The 14 neutralizing mAbs were characterized. **(A)** Percentage of mAbs capable of achieving 50% neutralization of SARS-CoV-2 pseudovirus at a concentration of 50 μg/ml from each donor **(B)** The IC_50_s of each neutralizing antibody in comparison to a negative control (AMMO1) are graphed. Each data point represents an independent replicate, and the bars indicate the mean. The non-RBD-binding mAbs, CV1-1 and CV3-25, left side of graph, are separated by a dashed line from the RBD-binding mAbs, right side of graph. **(C)** SARS-CoV-2 neutralizing mAbs were assessed for ability to neutralize SARS-CoV-1. CR3022 is a control SARS-CoV-1 neutralizing mAb. Full data in **Supplemental Figure 4 A-D**. **(D)** The IC_50_s of the inferred germline versions of the mAbs (open dots) are compared to IC_50_s of mutated mAbs (solid dots). Additional data in **Supplemental Figure 5**. **(E)** The area under the curve (AUC) of competition BLI for SARS-CoV-2 RBD is compared. Dots are shown as the median of two replicates with standard deviation indicated by error bars. The dotted line at the RBD-alone condition indicates BLI signal of uninhibited RBD:ACE2 binding. The NTD-specific CV1-1 mAb is used a negative control. mAbs that show competition a binding signal below the dotted line block ACE2 binding, mAbs with a binding signal above the dotted line enhance ACE2 binding by increasing avidity through immune complex formation. **(F)** Correlation between SARS-CoV-2 neutralization IC_50_ with area under the curve (AUC) of the BLI of competition with ACE2 for RBD binding. R^2^ value for nonlinear fit and Spearmen correlation p value are shown. **(G)** The area under the curve (AUC) of competition BLI for SARS-CoV-1 RBD is compared on this graph performed as in **E**. Full ACE2-competition data in **Supplemental Figure 4F-J**. Additional characterization of CV1-1 ad CV2-75 in **Supplemental Figure 6**.

**Figure 5:**
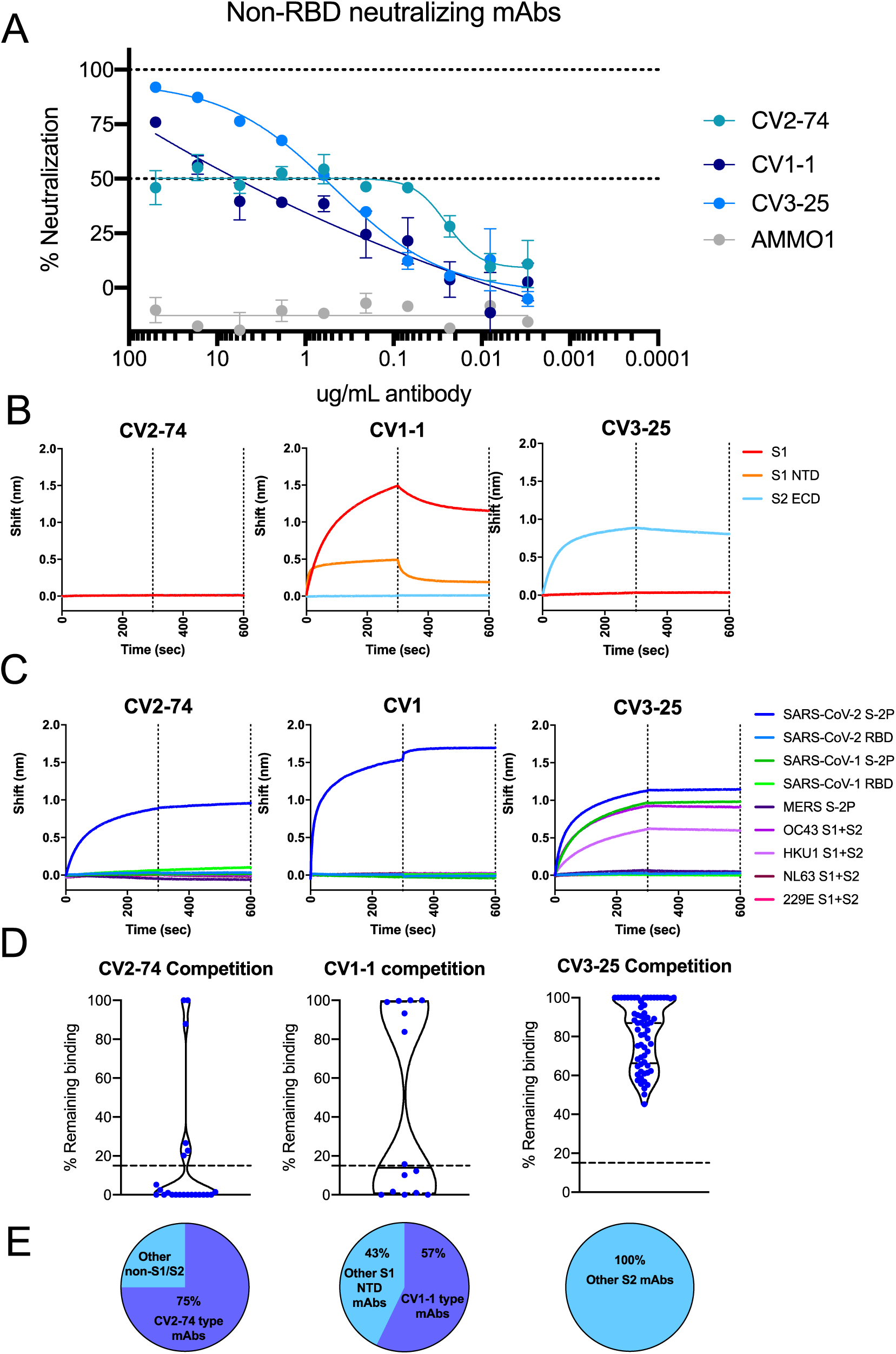
Neutralization by non-RBD-binding nAbs. The three neutralizing, non-RBD binding mAbs were characterized. **(A)** Neutralization curves for non-RBD binding mAbs. **(B)** BLI traces for the indicated mAbs binding to SARS-CoV-2 S1 or S2 subunits or the NTD subdomain of S1. **(C)** BLI traces of mAbs incubated with human coronavirus antigens as indicated. **(D)** Violin plots show competitions between each non-RBD mAb and other mAbs. Each data point represents the area under the curve (AUC) of an individual mAb binding to RBD (left), NTD (middle) or S2 (right) minus AUC of competition with either CV2-74 (left), CV1-1 (middle) or CV3-25 (right). Dotted line at 15% remaining binding indicates what is considered true competition, dots below the line are considered competitive. For CV1-1, S1 NTD mAbs from CV1, CV2 and CV3 were tested. For CV2-74, all non-S1/S2 mAb in all four sorts were tested. For CV3-25 all S2-binding mAbs in all four sorts were tested. Median of plot is indicated as solid line with quartiles indicated as dashed lines. **(E)** Pie charts show percentage of mAbs in each set that effectively compete with each tested mAb. mAbs that competed are indicated in the purple section while non-competitive mAbs are in blue.

The three most potent nAbs, all anti-RBD, were CV1-30 (IC_50_=0.044 μg/ml) (Seydoux et al., 2020), CV3-1 (IC_50_=0.007 μg/ml) and PCV19 (IC_50_=0.072 μg/ml). The anti-NTD mAb (CV1-1) had lower neutralizing potency (IC_50_=8.2 μg/ml) and as we previously reported (Seydoux et al., 2020), its maximum level of neutralization was lower than 100% **(Supplemental Figure 4C)**, similar to other anti-NTD mAbs (Liu et al., 2020). CV1-1 displayed decreased binding against more stable SARS-2-CoV S engineered proteins (S-6P), as shown by lower overall unit response and faster off-rate by BLI and does not bind like other published NTD-targeting antibodies by negative stain EM (**Supplemental Figure 5A,B**) (Liu et al., 2020). The IC_50_ of anti-S2 mAb, CV3-25, was 0.34 μg/ml, which is comparable to most anti-RBD nAbs with the exception of CV1-30, CV3-1 and PCV19.

Out of the 14 nAbs, seven (CV2-20, CV2-71, CV2-75, CV3-7, CV3-17, CV3-25 and CV3-43) also bound the S-2P of SARS-CoV-1 and four of the seven neutralized this virus **(Figure 4C, Supplemental Figure 4D and Supplemental Table 2)**. Three were anti-RBD (CV2-71, CV2-75 and CV3-17), while the fourth, CV3-25, bound to S2 **(Figure 5B, Supplemental Table 2)**. Interestingly, while the IC_50_s of CV2-71, CV2-75 and CV3-25 against SARS-CoV-1 and SARS-CoV-2 were not significantly different, CV3-17 neutralized SARS-CoV-1 more potently than SARS-CoV-2 **(Supplemental Figure 4E)**. Furthermore, the two most potent anti-SARS-CoV-2 mAbs (CV1-30 and CV3-1) did not neutralize SARS-CoV-1.

### Neutralization of inferred germline forms of mAbs

CV1-30 has only 2 non-silent somatic mutations (both in VH) that we previously reported are important for potent neutralization of SARS-CoV-2 (Hurlburt et al., 2020). To examine if this is a general phenomenon among anti-RBD SARS-CoV-2 nAbs, we generated the inferred-germline (iGL) versions of 6 anti-RBD Abs (CV2-20, CV2-71, CV2-75, CV3-1, CV3-7, and CV3-43) and measured their neutralizing potencies **(Figure 4D and Supplemental Fig 6A)**. Three of six anti-RBD iGL-nAbs, CV2-20 (3 amino acid mutations), CV2-75 (3 amino acid mutations) and CV3-43 (9 amino acid mutations) were non-neutralizing. However, no differences in neutralizing potency between the mutated and iGL-CV2-71 (3 amino acid mutations), iGL-CV3-1 (2 amino acid mutations) and iGL-CV3-7 (9 amino acid mutations) were observed. Reductions in neutralizing potency of the iGL mAbs correlated with faster dissociation rates from RBD **(Supplemental Fig 6B)**. The anti-NTD mAb CV1-1 has no amino acid mutations in its V genes while the anti-S2 Ab CV3-25 has 5 mutations. Reversion of the anti-S2 mAb CV3-25 to its germline form also led to a significant reduction in its neutralizing potency. Thus, some anti-SARS-CoV-2 nAbs are capable of potent neutralization in the absence of affinity maturation, while the neutralizing activity of others depends on the accumulation of a small number of mutations. Overall, however, there was no correlation between the neutralization potency and the degree of SHM (data not shown).

### Potent anti-RBD neutralizing antibodies block the binding of ACE-2 to RBD

We next examined whether the differences in neutralizing potencies of the anti-RBD nAbs **(Figure 4B)** were due to differences in their relative abilities to block the RBD-ACE2 interaction **(Figure 4E, Supplemental Figure 4F)**. While CV2-71, CV2-75 and CV3-1 abolished ACE2 binding to RBD, suggesting that they either directly bound the receptor binding motif (RBM) like CV1-30 (Hurlburt et al., 2020), or indirectly (sterically) hindered this binding, the remaining 7 anti-RBD NAbs (CV2-20, CV2-66, CV3-7, CV3-17, CV3-43, CV3-45 and PCV19) did not inhibit the RBD-ACE2 interaction. Similar observations were made when the abilities of mAbs to block the interaction of recombinant S-2P to cells expressing ACE2 were examined **(Supplemental Figure 4G)**. Indeed, a correlation between the potency of neutralization and the extent to which a mAb blocked the RBD-ACE2 interaction was observed **(Figure 4F and Supplemental Figure 4H)** in agreement with previous reports (Baum et al., 2020a; Brouwer et al., 2020; Gavor et al., 2020; Hoffmann et al., 2020; Wan et al., 2020a; Yan et al., 2020).

As mentioned above, three of the anti-RBD mAbs (CV2-71, CV2-75, and CV3-17) also neutralized SARS-CoV-1. The abilities of these antibodies to block the ACE2 interaction with the SARS-CoV-1 RBD were similar to their abilities to block the interaction of ACE2 with the SARS-CoV-2 RBD, with CV2-71 and CV2-75 blocking ACE2 interaction to some degree **(Figure 4G, Supplemental Figure 4I, J)**. Antibodies like CV1-30 and CV3-1 that potently neutralize SARS-CoV-2 and block the interaction between SARS-CoV-2 RBD and ACE2, fail to mediate SARS-CoV-1 neutralization because they bind the RBM in the RBD which has limited sequence homology to that of SARS-CoV-1 RBD (Hurlburt et al., 2020). In contracts, CV2-75 binds the RBD at an epitope distinct from the receptor binding motif (RBM) (**Supplemental Figure 5C, Supplemental Table 3**) and is only accessible when the RBD is in the up conformation. The residues that CV2-75 interacts with on the RBD are nearly completely conserved between SARS-CoV-1 and −2 explaining the cross-neutralizing ability (**Supplemental Figure 5D**). An alignment with the structure of ACE2-RBD, showed that the heavy chain of CV2-75 would clash with the glycan at Asn322 in ACE2, establishing a mechanism of competition (**Supplemental Figure 5E**).

### Neutralization by non-RBD binding Abs

As mentioned above CV2-74 binds to an undefined epitope on S, that is present on S-2P but absent or not properly presented on the recombinant S1 or S2 proteins used here **(Figure 5B)**. We identified several mAbs sharing this binding property (especially in PCV1) and the majority (75%) of these mAbs did compete the binding of CV2-74 to S-2P **(Figure 5D,E)**. The fact that among these mAbs only CV2-74 displayed neutralizing activity suggests that either the other mAbs bind distinct epitopes on S-2P and indirectly affect the binding of CV2-74 to S-2P or that CV2-74 binds a unique but overlapping epitope. It is note-worthy that CV2-74 displays an unusual neutralization curve, where the mAb neutralizes only 50% of the virus across a thousand-fold concentration range **(Figure 5A)**. For that reason, we did not assign an IC50 value to CV2-74.

Out of the 14 anti-NTD mAbs we identified, 8 (57%) competed the binding of CV1-1 to S-2P **(Figure 5D,E)** and yet, CV-1-1 was the only neutralizing anti-NTD mAb (**Figure 4B)**. Interestingly, CV1-1 displayed decreased binding to more stable SARS-CoV-2 engineered soluble proteins (**Supplemental Figure 5**). While BLI revealed binding of CV1-1 to recombinant NTD, the on-rate and maximal binding signal was lower than to the entire S1 domain, suggesting that secondary (or quaternary) contacts are important (**Figure 5B**). Indeed, negative-stain EM analysis indicates that it recognizes NTD differently than other anti-NTD mAbs (such as COVA1-22 (Brouwer et al., 2020), with a footprint that might also include an area just above the S1/S2 cleavage site (**Supplemental Figure 5B**).

Out of 87 anti-S2 mAbs, CV3-25 was the only one capable of neutralizing SARS-CoV-2 and SARS-CoV-1 (**Figure 4B and C**) and of binding the S proteins of the OC43 and HKU1 betacoronaviruses **(Figure 5C, Supplemental Table 2)**. As none of the other 86 anti-S2 mAbs competed the binding of CV3-25 to S2-P (**Figure 5D,E**) we expect that CV3-25 binds a unique epitope on the S2 subunit which is present not only on SARS-CoV-1 but also on the other coronaviruses tested here.

### Neutralizing mAbs as pre-exposure prophylaxis in k18-hACE2 mice

To assess whether nAbs with different epitope specificities offer the same level of protection *in vivo,* we compared the protective abilities of CV1-1, CV1-30 and CV2-75 in the K18-hACE2 mouse model (Winkler et al., 2020). As discussed above, CV1-1 binds NTD and has an IC50 of 8.2 μg/ml, while CV1-30 and CV2-75 bind the RBD and have IC50s of 0.044 and 1.7 μg/ml, respectively. Thus, CV1-1 and CV2-75 have similar neutralizing potentials but recognize different regions of the viral spike.

Mice were given a dose of 10 mg/kg of CV1-1, CV2-75, CV1-30, or an isotype control anti-EBV antibody AMMO1 (Snijder et al., 2018), and then challenged intranasally with 10,000 plaque forming units (PFU) of SARS-CoV-2 (**Figure 6A**). Two days post challenge, half of the animals were euthanized to assess viral loads in the lung, and the remaining 5 animals were monitored for survival for up to 14 days. Two days post-challenge, mice receiving AMMO1, CV1-1 and CV2-75 had high levels (1 x 10^8^ PFU) of infectious virus and viral RNA in the lung (**Figure 6B and C**). 3 of 5 remaining animals in the CV1-1- and CV2-75 groups did not survive beyond 6 days post-challenge (**Figure 6C,D**). In contrast, CV1-30 significantly limited viral replication in the lungs at 2 days post-challenge (**Figure 6B,C**) and all remaining mice survived (**Figure 6D**).

**Figure 6.**
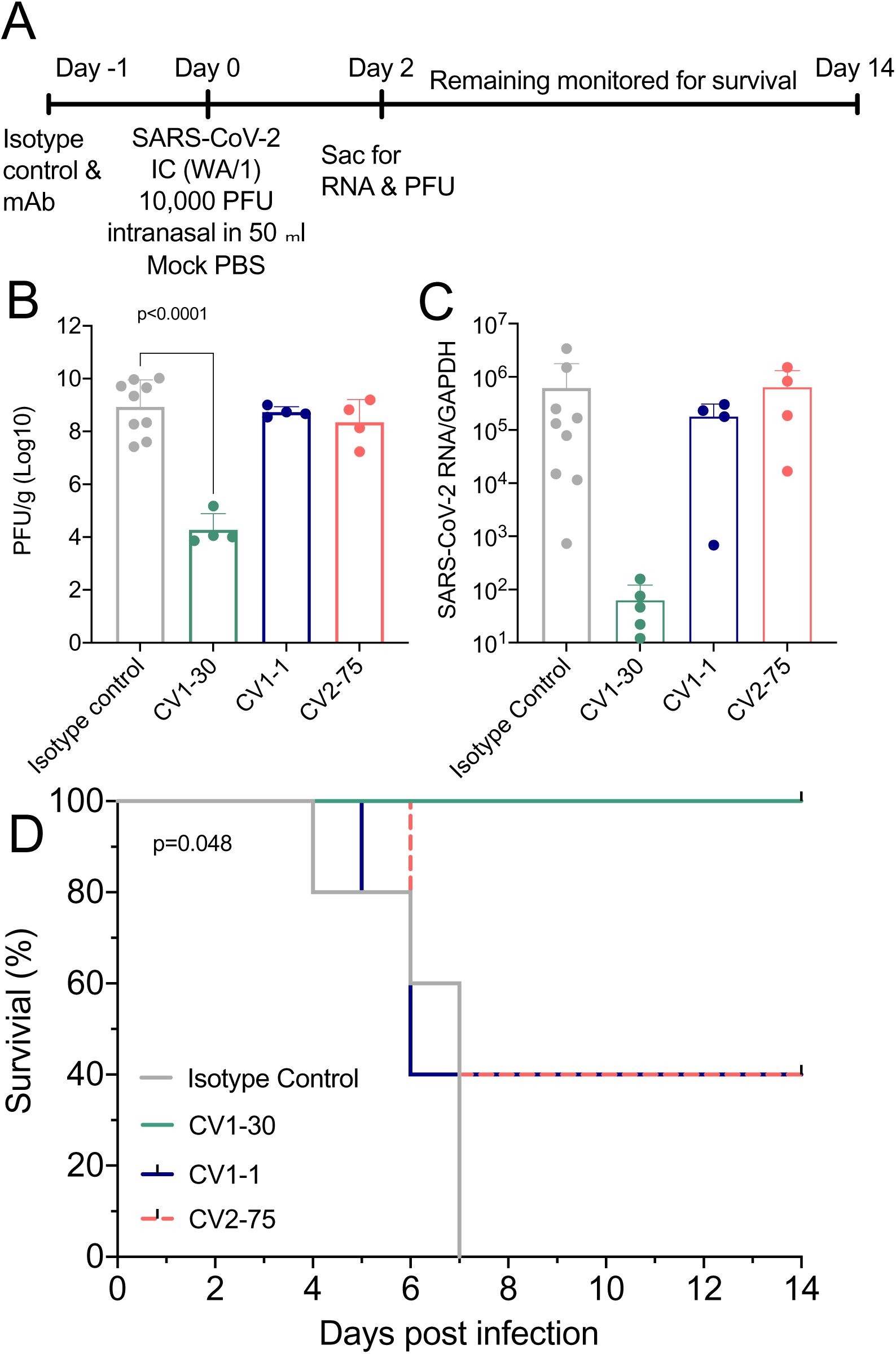
Neutralizing mAbs as pre-exposure prophylaxis in k18-hACE2 mice. CV1-1, CV1-30 and CV2-75 were assessed to see whether they could confer protection in a mouse model. **(A)** Experimental timeline. **(B)** Number of plaque-forming units in the lungs 2 days following challenge. **(C)** viral RNA in lung tissue 2 days after challenge was measured by qPCR and normalized to GAPDH expression. **(D)** Kaplan-Meyer survival curve of the viral load/titer in the lungs of remaining mice comparing the various treatment groups. Statistics determine by One-Way ANOVA with Dunnett’s multiple comparison test. *p<0.05, **p<0.01, ***p<0.001, ****p<0.0001

Collectively these data indicate that neutralizing potency rather than epitope specificity is the most important factor in defining the prophylactic efficacy of anti-SARS-CoV-2 antibodies. Consistent with this, CV3-25 also showed partial protection in a K18-hACE2 animal model (see accompanying manuscript: by Ullah et al., *“Live imaging of SARS-CoV-2 infection in mice reveals neutralizing antibodies require Fc function for optimal efficacy”*).

### Neutralization of the mutant SARS-CoV-2 B.1.351 variant

Recently, lineages of viral variants have emerged in the United Kingdom (B.1.1.7), South Africa (B.1.351), and Brazil (P.1) that harbor specific mutations in their S proteins that may be associated with increased transmissibility (Davies et al., 2020; Faria et al., 2021; Rambaut et al., 2020; Sabino et al., 2021; Tegally et al., 2020; Volz et al., 2021). The B.1.351 lineage appears to be more resistant to convalescent sera and mAbs (Edara et al., 2021; Liu et al., 2021; Stamatatos et al., 2021; Wang et al., 2020b; Wibmer et al., 2021; Wu et al., 2021). It is defined by several mutations in RBD (K417N, E484K, N501Y), NTD (D80A, D215G,) and S2 (D614G) (O’Toole et al., 2021; Tegally et al., 2020). Other mutations are also found in the B.1.351 lineage in the NTD R246I and deletion 242-244, and S2 A701V, but at lower frequencies.

We recently reported that these mutations abrogated the neutralizing activity of CV1-1 and reduced the neutralizing activities of the two most potent nAbs CV1-30 and CV3-1 (Stamatatos et al., 2021). Here we evaluated the ability of the 4 cross-neutralizing mAbs (CV2-75, CV3-17, CV2-71 and CV3-25) to neutralize the B.1.351 mutant strain **(Figure 7)**. We found that all four mAbs retained their neutralizing activities against B.1.351.

**Figure 7.**
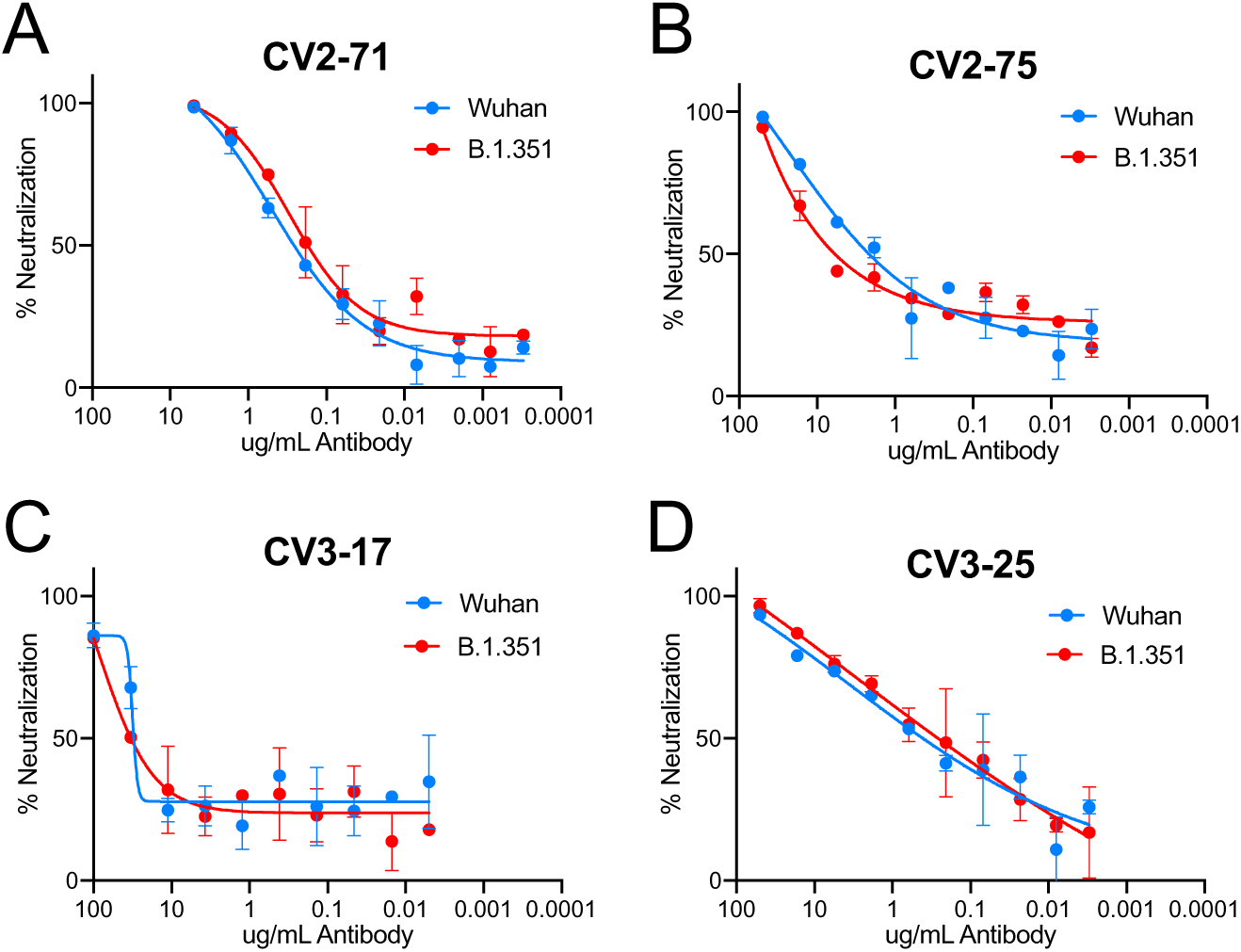
Neutralization of the mutant B.1.351 variant. The SARS-CoV-1 neutralizing mAbs were tested against the B.1.351 strain. **(A)** CV2-71. **(B)** CV2-75. **(C)** CV3-17. **(D)** CV3-25. Graphs show neutralization curves for the Wuhan strain of SARS-CoV-2 in blue and the curve for the B.1.351 strain in red.

## DISCUSSION

Our study reveals that naïve B cells expressing VH3-30 and VH1-18 preferentially recognize the SARS-CoV-2 envelope spike, but that nAbs are produced by B cells expressing diverse BCRs. Of the 198 mAbs characterized here (isolated at 3-7 weeks post-symptom development), 14 (7%) displayed neutralizing activities and among them, only CV3-7 was derived from VH3-30. In fact, the 11 anti-RBD nAbs were derived from distinct B cell clones, that cross-competed for binding, and 4 prevented the RBD-ACE2 interaction. These observations, combined with the fact that anti-RBD nAbs can neutralize the virus with no, or minimal somatic mutation, may explain why potent anti-SARS-CoV-2 neutralizing antibody responses are rapidly generated within a few weeks of infection, or shortly following 2 immunizations with vaccines that express the viral spike (Jackson et al., 2020; Walsh et al., 2020). The observation that 7 of 11 anti-RBD nAbs do not prevent the RBD-ACE2 interaction, indicates different mechanisms of neutralization by anti-RBD antibodies. The former nAbs may prevent RBD-heparin interactions (Clausen et al., 2020), stabilize the RBDs in their ‘up’ conformation and thus prematurely activate the fusion machinery (Koenig et al., 2021; Wrapp et al., 2020a), or limit the conformational changes, and particularly the RBD movement, that are required for cell fusion, allowing them to neutralize without directly blocking ACE2 binding.

The two most potent anti-SARS-CoV-2 nAbs, CV1-30 and CV3-1, which both bind SARS-CoV-2 RBD but not SARS-CoV-1 RBD, did not neutralize SARS-CoV-1 while CV2-75 and CV3-17, which bind not only SARS-CoV-2 RBD but also SARS-CoV-1 RBD and displayed weaker anti-SARS-CoV-2 neutralizing activities, were able to efficiently neutralize SARS-CoV-1. A comparison of the CV2-75-RBD and CV1-30-RBD (Hurlburt et al., 2020) structures reveal that CV2-75 binds an area of SARS-CoV-2 RBD with higher sequence homology with SARS-CoV-1 RBD. In contrast, CV1-30 binds directly to the receptor binding motif which only has 50% sequence homology among SARS-CoV-1 and SARS-CoV-2 (Finkelstein et al., 2021; Hurlburt et al., 2020; Wan et al., 2020b).

The mechanisms of neutralization of the three non-RBD binding nAbs characterized here (CV1-1, CV2-74 and CV3-25) are presently unknown. As CV1-1, CV2-74 and CV3-25 do not interfere with the binding of ACE-2 to S-2P, we anticipate that they mediate neutralization by interfering with a step in the fusion process that follows attachment. The viral spike undergoes conformational changes, specifically in the S2 region, during virus-cell binding and fusion (Cai et al., 2020; Gavor et al., 2020; Walls et al., 2020). Potentially, these three mAbs may prevent these conformational changes from occurring, either by locking the spike in an intermediate conformation, preventing cleavage or by stabilizing its pre-fusion conformation.

The fact that the binding of CV1-1 to S-2P was competed by the other anti-NTD mAbs (14 total), all of which were non-neutralizing, suggests that CV1-1 recognizes the NTD in a distinct manner from the non-neutralizing anti-NTD mAbs. Similarly, the binding of CV2-74 to S-2P was competed by the other non-neutralizing non-S1/S2 mAbs (23 total), which strongly suggests that these mAbs all recognize the same immunogenic region, but that CV2-74 recognizes it in a unique manner. In contrast, none of the anti-S2 mAbs isolated here (65 total) competed the binding of CV3-25 to S-2P. These observations and the fact that CV3-25 potently neutralizes both SARS-CoV-1 (IC_50_ 2.1 μg/mL) and SARS-CoV-2 (IC_50_ 0.34 μg/mL) and the B.1.351 mutant strain and binds the S proteins of HKU1 and OC43, strongly suggests that it recognizes a conserved epitope among diverse coronaviruses. As only two other anti-S2 antibodies that neutralize both SARS-CoV-1 and SARS-CoV-2, but with weaker neutralizing activities than CV3-25, were reported so far (Song et al., 2020; Wang et al., 2020b) we expect the epitope of CV3-25 to be less immunogenic than those recognize by non-neutralizing anti-S2 antibodies.

We propose that because of its cross-neutralizing activity, its ability to neutralize the B.1.351 and because it binds the OC43 and HKU1 spikes, CV3-25 represents an antibody type that a pan-coronavirus vaccine should elicit. We expect that the protective potentials of such antibodies will improve through the accumulation of amino acid mutations in their VH and VLs through sequential immunizations. As a first step, the epitope of CV3-25 must be identified, and immunogens should be designed expressing it in the most immunogenic form.

In summary, our study indicates that neutralization of SARS-CoV-2 and SARS-CoV-1 does not necessitate the expansion of B cell lineages that express particular VH/VL pairings and that even the unmutated forms of some antibodies can potently neutralize SARS-CoV-2 and SARS-CoV-1. As these viruses are capable of tolerating mutations in distinct regions of its viral spike, they will be able to escape the neutralizing activities of most nAbs. The S2 subunit, however, contains at least one epitope that although poorly immunogenic, is present on four of five human beta coronaviruses. That epitope, as defined by its recognition by CV3-25 is a valid candidate for the development of a global coronavirus vaccine.

## AKNOWLEDGEMENTS

This work was supported by generous donations to the Fred Hutch COVID-19 Research Fund, by grants (P51 OD011132 and 3U19AI057266-17S1) from the National Institute of Allergy and Infectious Diseases (NIAID), National Institutes of Health (NIH), by the Bill and Melinda Gates Foundation (OPP1170236/INV-004923), by the Emory Executive Vice President for Health Affairs Synergy Fund award, the Pediatric Research Alliance Center for Childhood Infections and Vaccines and Children’s Healthcare of Atlanta, the Woodruff Health Sciences Center 2020 COVID-19 CURE Award and the Emergent Ventures Award (HYC). Support was also provided by le Ministère de l’Économie et de l’Innovation du Québec, Programme de soutien aux organismes de recherche et d’innovation to A.F., by the Fondation du CHUM and by a CIHR foundation grant #352417 to A.F. A.F. is the recipient of Canada Research Chair on Retroviral Entry no. RCHS0235 950-232424. We thank the J. B. Pendleton Charitable Trust for its generous support of Formulatrix robotic instruments. Results shown in this report are derived from work performed at Argonne National Laboratory, Structural Biology Center (SBC), ID-19, at the Advanced Photon Source. SBC-CAT is operated by U Chicago Argonne, LLC, for the U.S. Department of Energy, Office of Biological and Environmental Research under contract DE-AC02-06CH11357. We also would like to thank L. Kehoe, S. P. Canny, K. Nanda and J. Czartoski for the care and enrollment of patients.

## AUTHOR CONTRIBUTIONS

<u>Conceptualization</u>, M.F.J., M.S.S., A.T.M, M.P., and L.S.; <u>Investigation</u>, M.F.J., A.J.M, N.R.A., J.F., L.J.H., N.K.H., E.S., Y.W, A.B.S., V.V.E., K.F., R.P., A.B.W., G.O., J.L.T., N.DR, E.S.Y., R.E.W, K.W.C, M.P; <u>Resources</u>, J.R.M., H.Y.C., J.A.E., M.S., M.J.M. and A.F.; <u>Writing</u>: M.F.J., A.T.M., and L.S.; <u>Supervision</u>, A.T.M., and L.S., Funding acquisition: A.T.M., L.S, M.S and M.J.M.

---

**Supplemental Figure 1:**
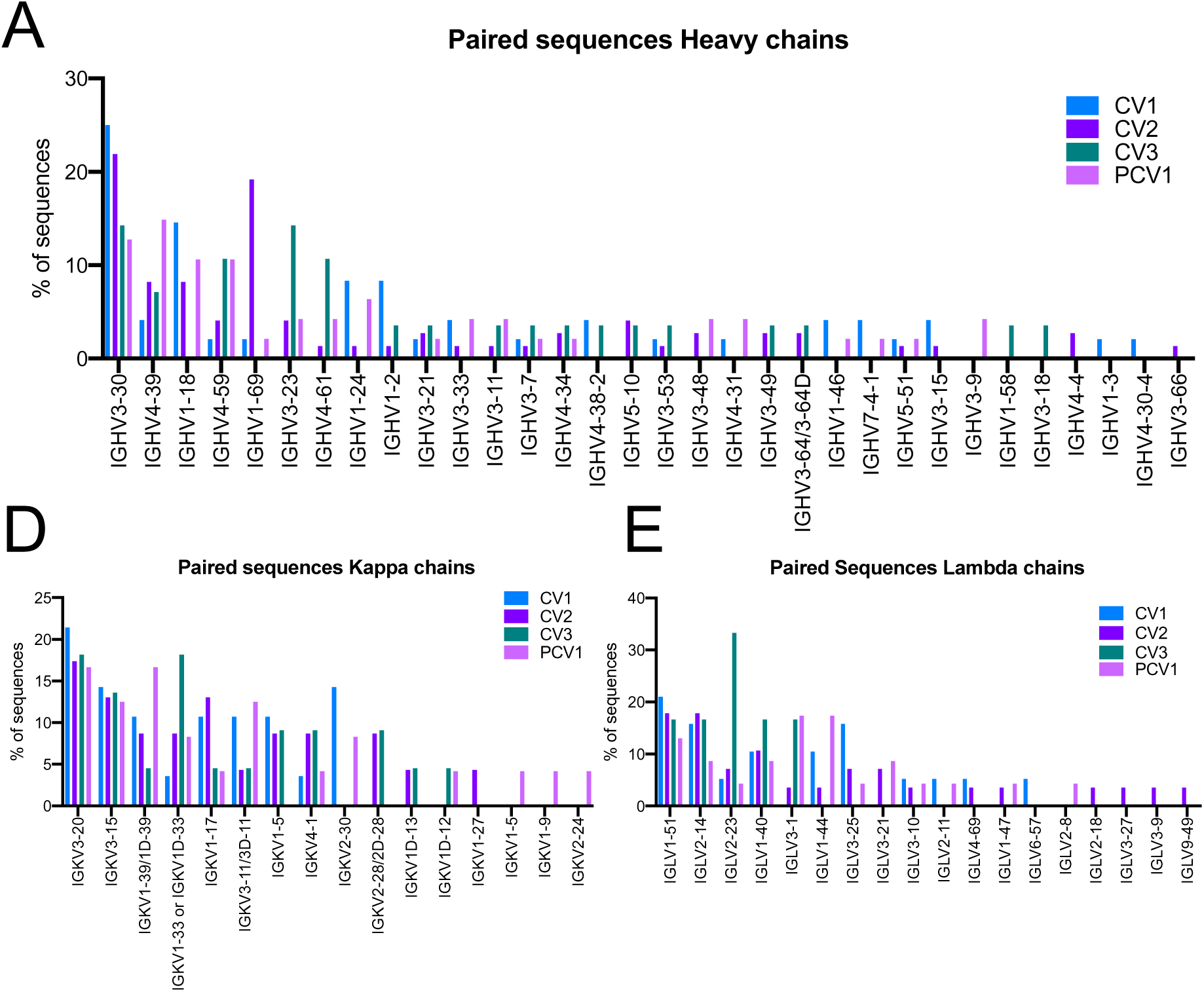
SARS-CoV-2 mAb VH and VL sequencing. Related to Figure 2. Full sequencing data for all VH and VL genes isolated from the four SARS-CoV-2 positive patients. **(A-C)** Full gene analysis for all paired heavy **(A)**, Kappa **(B)**, and Lambda **(C)** sequences from all four sorts displayed as above as percentage of total sequences from each sort.

**Supplemental Figure 2:**
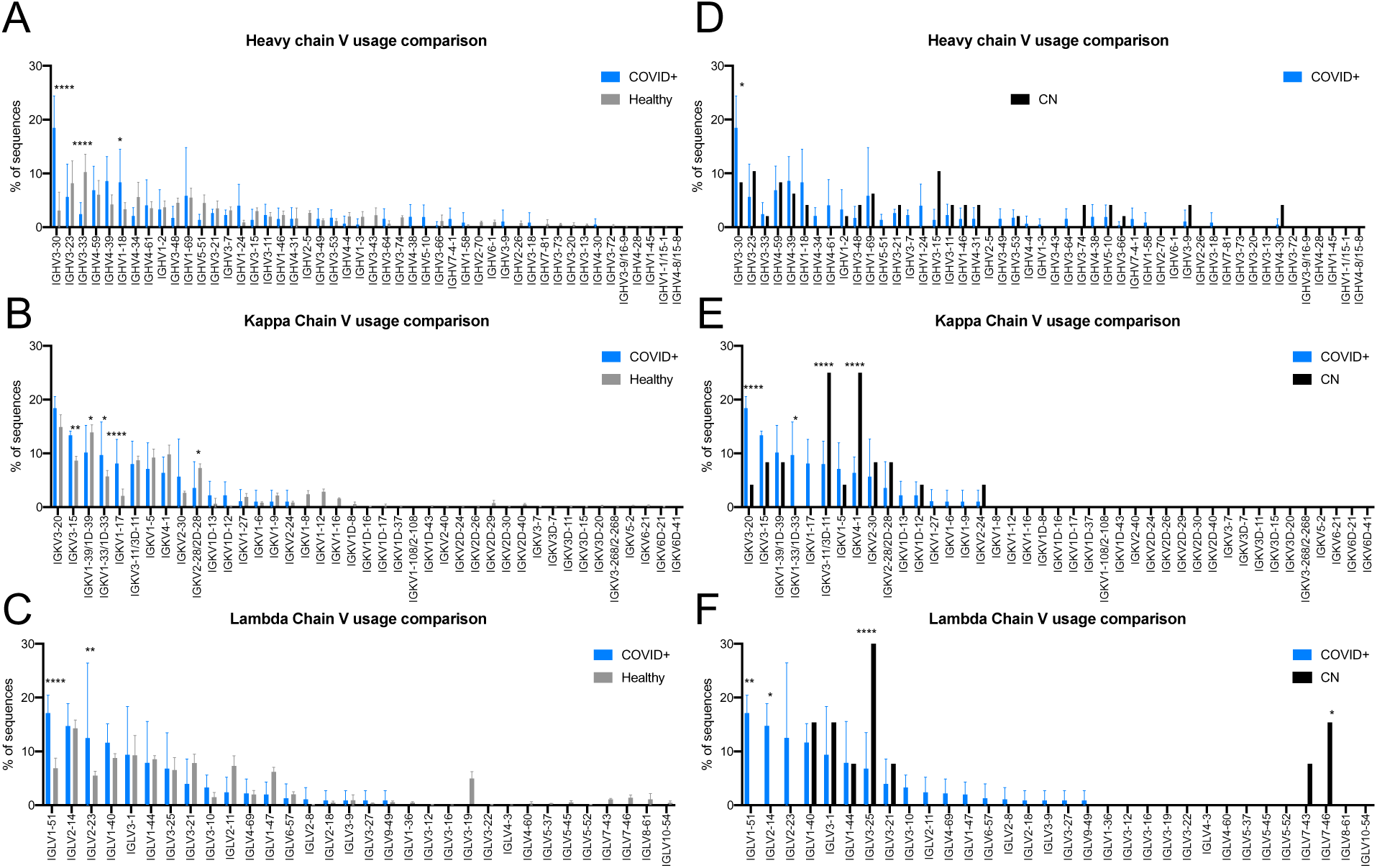
Comparison of VH and VL usage to healthy and naïve repertoire. Related to Figure 2. The full VH and VL usage in the four SARS-CoV-2 patients was compared to the usage in unbiased sequencing of health individuals and to VH and VL usage by the S-2P+ CN sort. **(A-C)** Comparison of V gene frequencies expressed in total B cells in the 5 10X sorted unexposed individuals to spike specific B cells from COVID+ donors. COVID+ sample V gene frequency is indicated in blue bars while healthy samples are indicated in grey bars for heavy **(A)**, kappa **(B)**, and Lambda **(C)** genes. **(D-F)** Comparison of V gene frequencies from S-2P+ B cells in CN, the naïve individual to spike specific B cells from COVID+ donors. COVID+ sample V gene frequency is indicated in blue bars while naive samples are indicated in black bars for heavy **(D)**, kappa **(E)**, and Lambda **(F)** genes. Significance calculated using two-way-ANOVA. Statistics between different samples evaluated as two-way-ANOVA with Šídák’s multiple comparison test. *p<0.05, **p<0.01, ***p<0.001, ****p<0.0001.

**Supplemental Figure 3:**
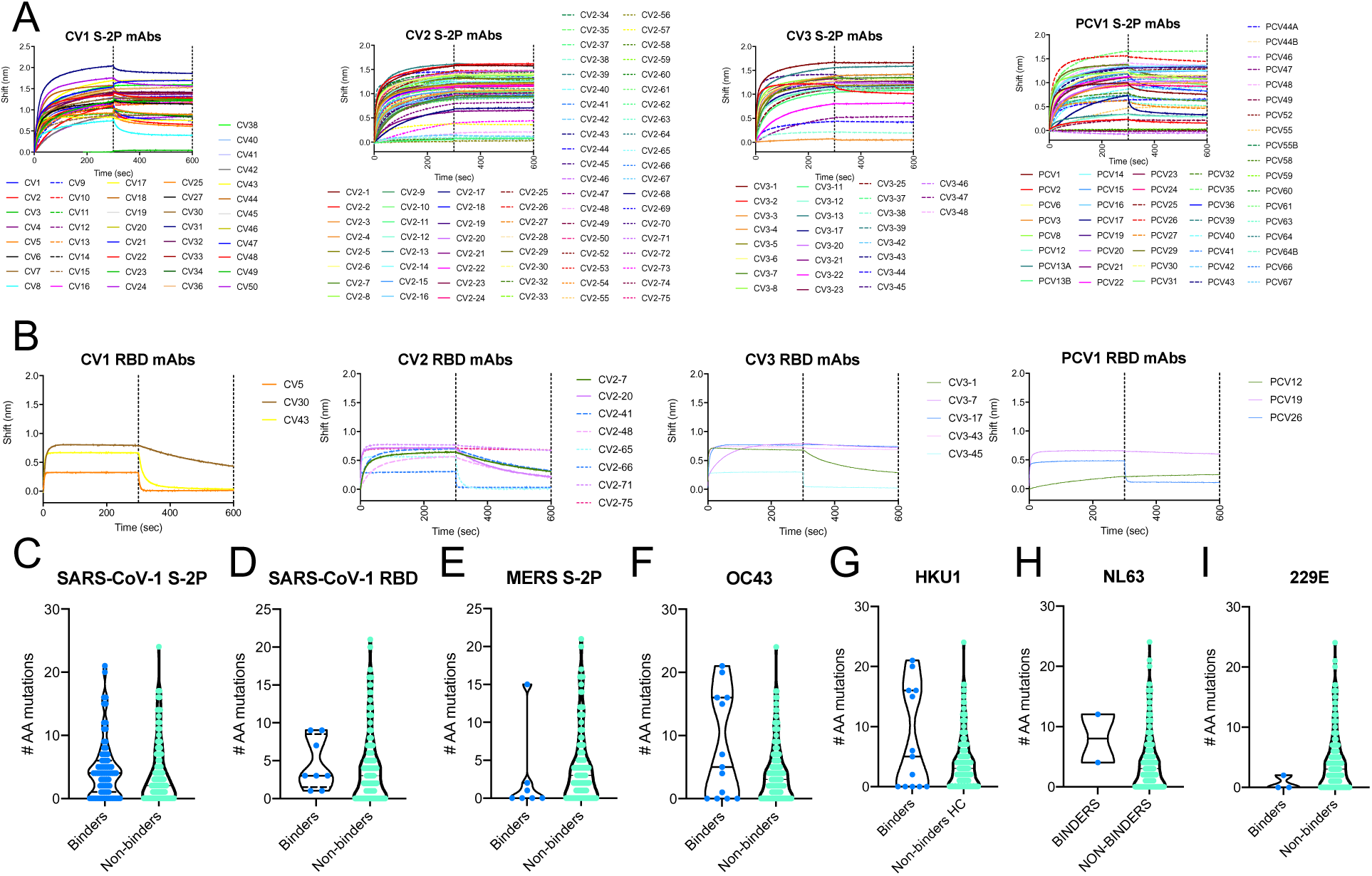
mAb epitope mapping and cross-reactivity BLI. Related to Figure 3. The binding of the 198 SARS-CoV-2-specific mAbs was assessed by BLI for epitope and cross-reactivity. **(A, B)** All spike-specific mAbs isolated from COVID+ donors were tested by BLI for binding to SARS-CoV-2 S2P **(A)** and SARS-CoV-2 RBD **(B). (C-H)** The number of amino acid mutations (sum of heavy chain and light chain mutations) for each for each SARS-CoV-2 mAb that bound (blue dots) or didn’t bind (teal dots) SARS-COV-1 S-2P **(C),** SARS-COV-1 RBD **(D)**, MERS S-2P **(E)**, OC43 **(F)**, HKU1 **(G)**, NL63 **(H)**, or 229E **(EI)**. Statistics were assessed by Mann-Whitney test, no comparisons were significant.

**Supplemental Figure 4:**
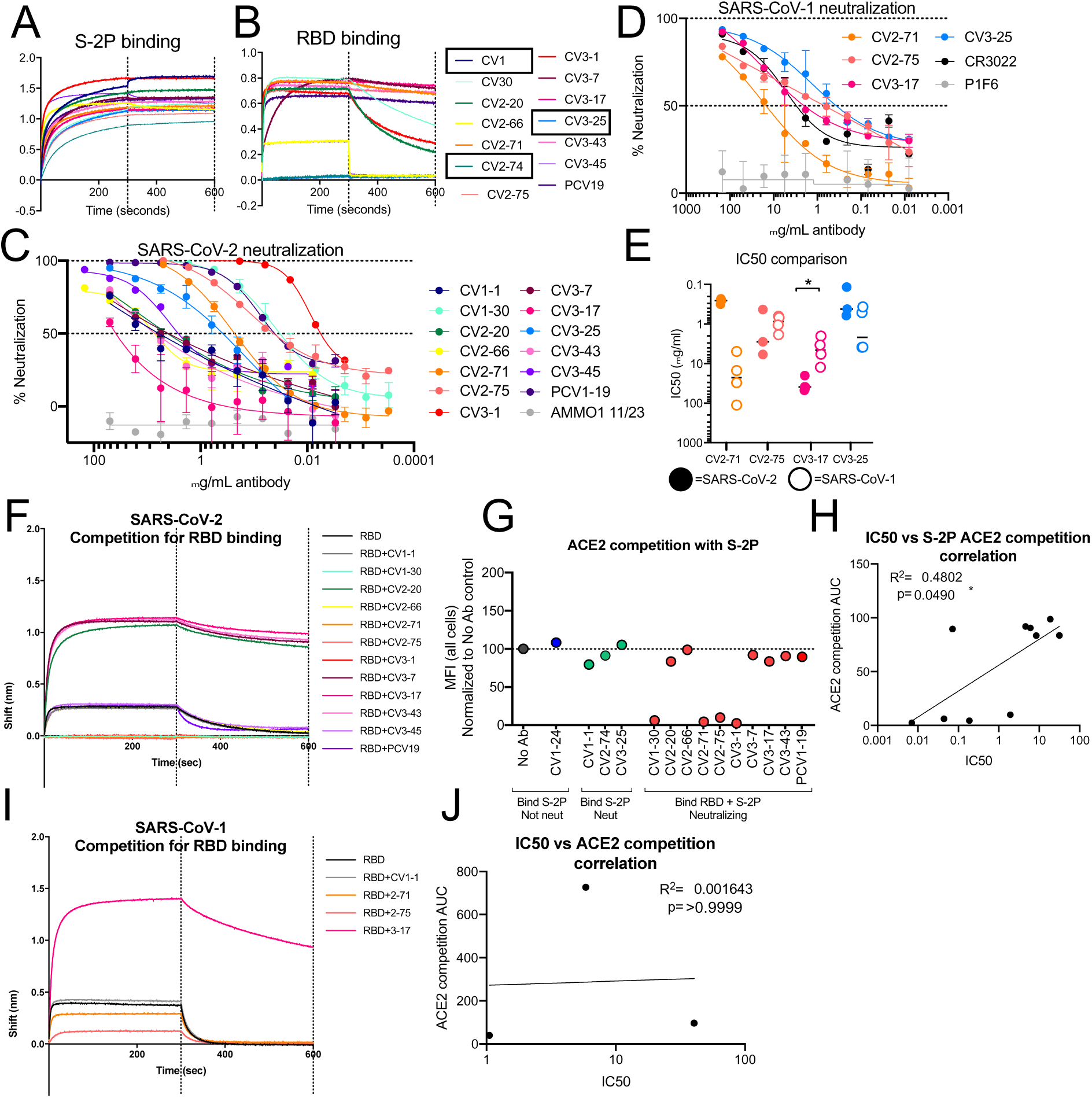
mAb Neutralizing potential and ACE2 competition. Related to Figure 4. **(A)** Binding of nAbs to SARS-CoV-2 S-2P as measured by BLI**. (B)** Binding of nAbs SARS-CoV-2 RBD as measured by BLI. Black boxes indicate non-RBD binding nAbs **(C)** Representative neutralization curves for all SARS-CoV-2 nAbs. **(D)** Representative neutralization curves for the indicated mAbs vs SARS-CoV-1. **(E)** Comparison of IC_50_s for mAbs that bind both SARS-CoV-2 and SARS-CoV-1. Each dot represents an independent replicate. Solid dots represent IC_50_s for SARS-CoV-2 neutralization while open dots represent IC_50_ for SARS-CoV-1 neutralization. Statistics evaluated by mixed-effect analysis. *p<0.05. **(F)** Competition between ACE2 and the indicated mAbs for SARS-CoV-2 RBD binding was measured by BLI. **(G)** Inhibition of fluorescently labeled S-2P binding to ACE2 expressing cells by the indicated nAbs was performed by flow cytometry. The dotted line indicates the MFI of S-2P binding in the absence of mAb. mAbs with values below this line show blocking of ACE2 binding to S-2P. **(H)** Correlation between mAb neutralization IC_50_ and area under the curve (AUC) of competition between mAb and ACE2 for S2P binding. R^2^ value for nonlinear fit and Spearmen correlation p value are shown. **(I)** Competition between ACE2 and the indicated mAbs for SARS-CoV-1 RBD binding was measured by BLI. **(J)** Correlation between mAb neutralization IC_50_ and area under the curve (AUC) of competition between mAb and ACE2 for SARS-CoV-1 RBD binding. R^2^ value for nonlinear fit and Spearmen correlation p value are shown on graph.

**Supplementary Figure 5:**
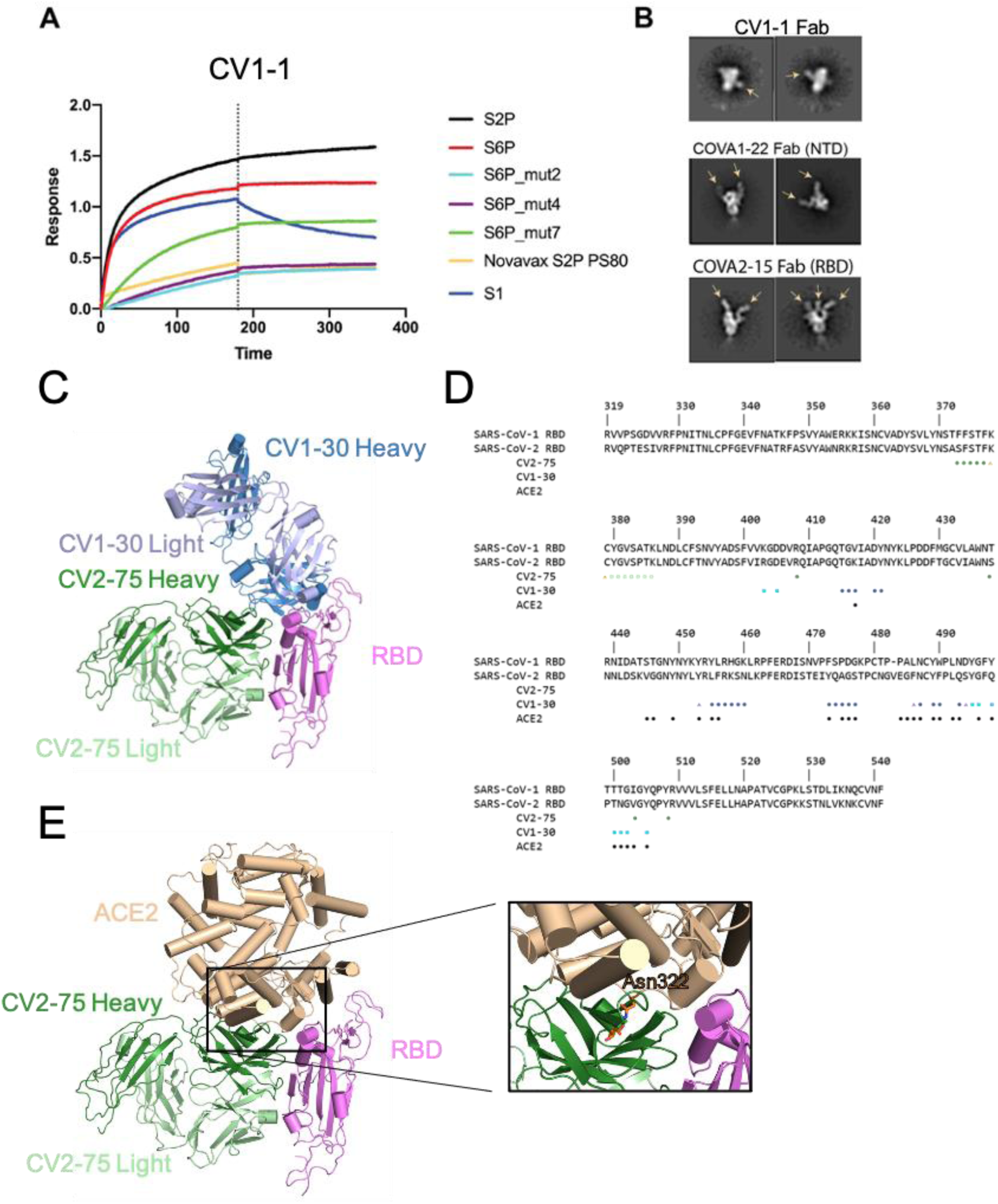
CV1-1 and CV2-75 characterization. Related to Figure 4 and 5. **(A)** Biolayer interferometry of immobilized CV1 IgG dipped into recombinant variants of SARS-2-CoV Spike. Vertical dotted line represents transition from association to dissociation steps. 2P = two proline stabilizing mutations, 6P = six proline stabilizing mutations. Mut2, mut4 and mut7 represent additional stabilizing mutations in S1 and/or S2. The Novavax S2P is formulated with polysorbate 80 and forms nanoparticles (Bangaru et al., 2020). **(B)** Representative EM 2D class averages from negatively-stained complexes of Fab and recombinant S protein. Arrows indicate Fab densities. COVA1-22 and COVA2-15 represent canonical NTD and RBD targeting antibodies, respectively (Brouwer et al., 2020). **(C,D)** Structural characterization of CV2-75 Fab bound to RBD indicates binding to a cryptic epitope. **(C)**. Cartoon representation of CV2-75 Fab (green) bound to RBD (pink) with structure of CV1-30 Fab (blue, PDB ID: 6XE1) superimposed. CV2-75 binds an epitope present only in the “up” RBD conformation. **(D)**. SARS-CoV-1 and SARS-CoV-2 RBD sequence alignment indicates that CV2-75 Fab to a conserved region between the two strains. Circles show heavy chain interactions to RBD, squares show light chain interaction, and triangles show both chains. **(E)**. RBD-ACE2 (PDB ID: 6M17) superposition to RBD-CV2-75 indicates clashes with glycans at Asn332 of ACE2.

**Supplemental Figure 6:**
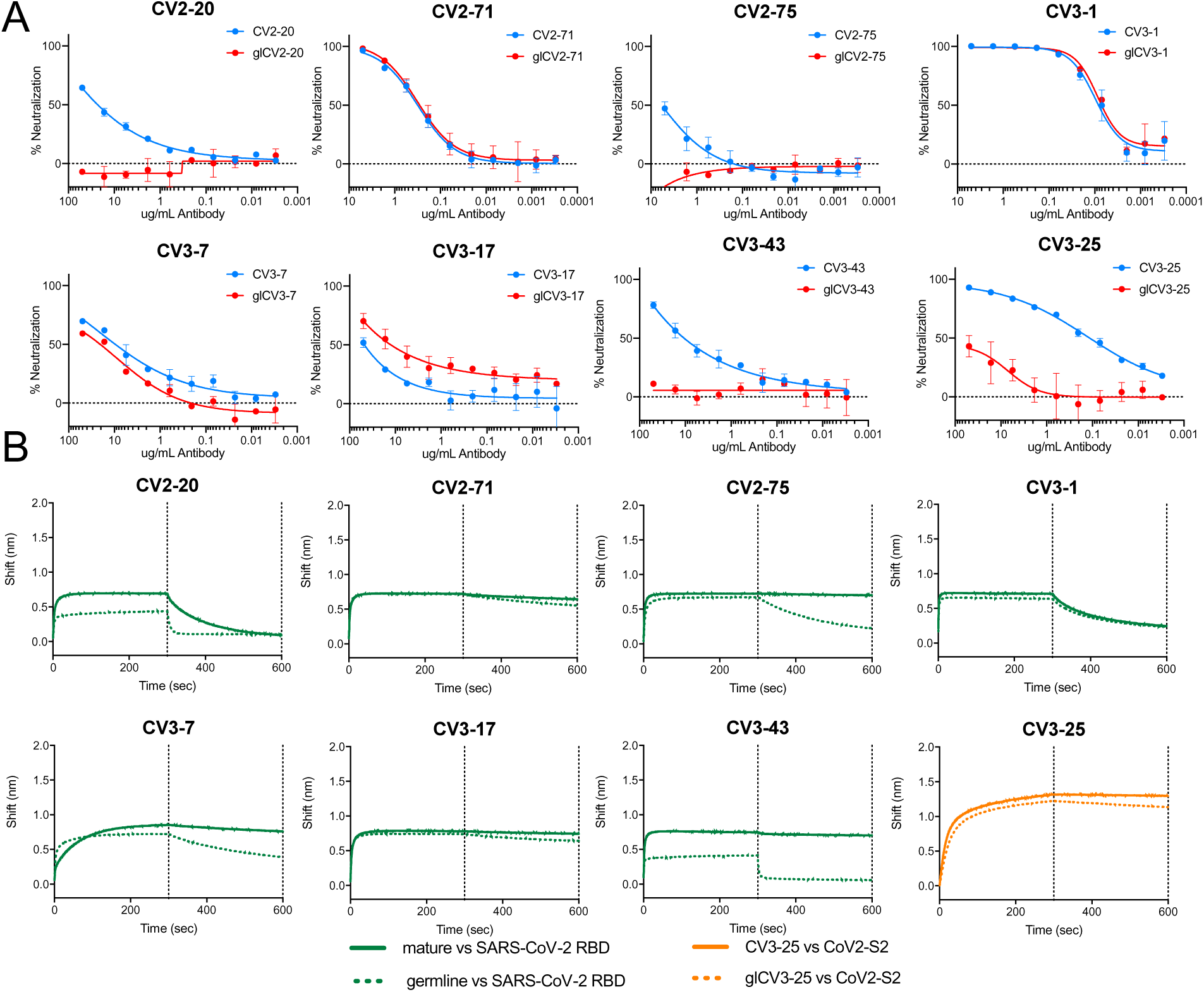
Neutralization potential of Inferred germline versions of mAb. Related to Figure 4. Versions of Nabs reverted to their germline forms were created and tested for neutralization potential and ability to bind their epitope. **(A)** Neutralization curves for mature (blue) and inferred germline (teal) mAbs. **(B)** The binding of the mature (solid lines) and inferred germline versions of nAbs (dotted lines) to the indicated antigens was measured by BLI. Binding to SARS-CoV-2 S2P (blue) was compared. For CV3-25, binding to SARS-Cov-1 S2 (orange) is shown.

**Supplemental Table 1:** B cell sorts from COVID-19 patients. Related to Figure 1 and 2.

| Donor | Weeks post symptom onset | PBMCs sorted | B cells sorted | %S-2P++ out of total B cells | %RBD++ out of total B cells | %RBD+ out of S-2P++ B cells | Unpaired Heavy chains | Unpaired Kappa chains | Unpaired Lambda chains | mAbs produced |
| --- | --- | --- | --- | --- | --- | --- | --- | --- | --- | --- |
| Patient 1 (CV1) | 3 | ~18 million | 768 | 0.65 | 0.0352 | 5 | 103 | 97 | 90 | 48 |
| Patient 2 (CV2) | 3.5 | ~10 million | 384 | 0.37 | 0.0482 | 12.7 | 132 | 125 | 112 | 73 |
| Patient 3 (CV3) | 6 | ~10 million | 432 | 0.23 | 0.0265 | 13 | 38 | 98 | 70 | 27 |
| Patient 4 (PCV1) | 7 | ~10 million | 1736 | 1.84 | 0.174 | 9.4 | 68 | 33 | 31 | 46 |
| Healthy sort (CN)<sub>a</sub> | N/A | ~5.7/~5.8 million | 70/96 | 0.104/0.128 | 0.015/0.019 | 11.3/9 | 24/42 | 37/24 | 15/15 | 9/28 |

a. The healthy sort, CN represents two distinct sorts of the same healthy individual. Values for each individual sort are separated by a forward slash in the table

**Supplemental Table 2:**
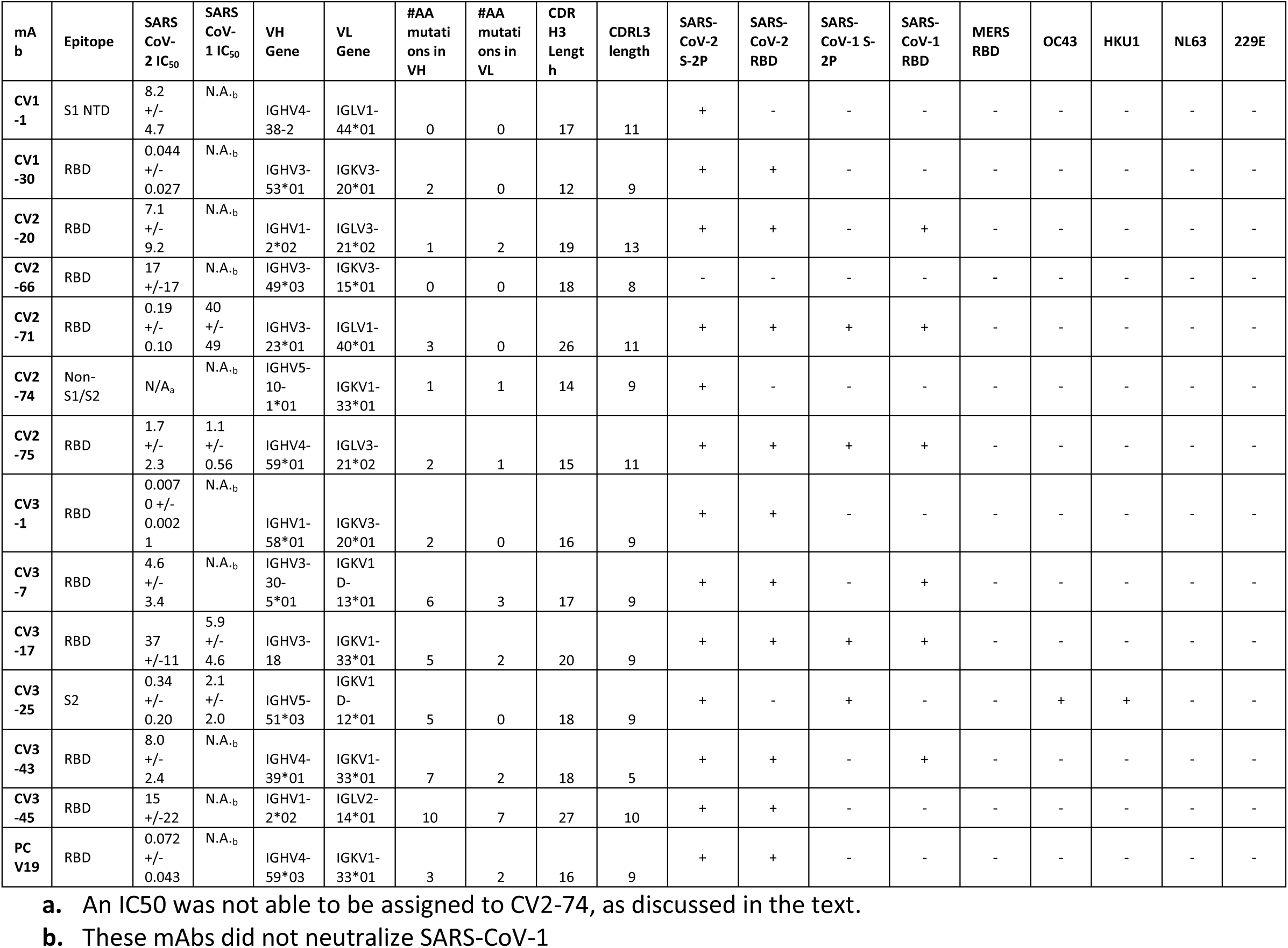
SARS-CoV-2 neutralizing mAbs. Related to Figure 3 and 4. This table shows the 14 neutralizing mAbs isolated and their binding epitopes, neutralization IC50, the VH and VL genes they are derived from, the number of mutations in these genes, the length of their CDRH3 and CDRL3, and whether they bind to SARS-CoV-2, SARS-CoV-1 and the other endemic human coronaviruses.

**Supplemental Data Table 3.** Data collection and refinement statistics for crystal structure. **Related to Figure 4 and Supplemental Figure 5**.

| CV2-75 Fab with SARS-CoV-2 RBD |  |
| --- | --- |
| **Data collection** |  |
| Space group | P2<sub>1</sub>2<sub>1</sub>2<sub>1</sub> |
| Cell dimensions |  |
| *a*, *b*, *c* (Å) | 86.37, 127.52, 155.67 |
| $\alpha$, $\beta$, $\gamma$ (°) | 90, 90, 90 |
| Resolution (Å) | 49.32-2.80 (2.90 – 2.80) |
| *R*<sub>merge</sub><sup>a</sup> | 0.034 (0.612) |
| <\|*I*/σ(*I*)> | 12.10 (1.44) |
| CC<sub>1/2</sub> | 0.999 (0.566) |
| Completeness | 99.48 (98.13) |
| Redundancy | 2.0 (1.99) |
| **Refinement** |  |
| Resolution (Å) | 49.32 – 2.80 (2.90 – 2.80) |
| No. unique reflections | 42798 (4141) |
| *R*<sub>work</sub><sup>b</sup>/*R*<sub>free</sub><sup>c</sup> | 24.52/28.11 (36.32/40.74) |
| No. atoms | 9874 |
| Protein | 9830 |
| Water | 16 |
| Ligand | 28 |
| B-factors (Å<sup>2</sup>) | 101.12 |
| Protein | 101.19 |
| Water | 59.90 |
| Ligand | 101.33 |
| RMS bond length (Å) | 0.003 |
| RMS bond angle (°) | 0.65 |
| **Ramachadran Plot Statistics<sup>d</sup>** |  |
| Residues | 1284 |
| Most Favored region | 92.89 |
| Allowed Region | 6.32 |
| Disallowed Region | 0.79 |
| Clashscore | 5.06 |
| **PDB ID** | **7M3I** |

<sup>a</sup> $R_{\text{merge}} = [\sum_h \sum_i |I_h - \bar{I}_{hi}| / \sum_h \sum_i I_{hi}]$ where $\bar{I}_h$ is the mean of $I_{hi}$ observations of reflection $h$ . Numbers in parenthesis represent highest resolution shell. <sup>b</sup> $R_{\text{factor}}$ and <sup>c</sup> $R_{\text{free}} = \sum ||F_{\text{obs}}| - |F_{\text{calc}}|| / \sum |F_{\text{obs}}| \times 100$ for 95% of recorded data ( $R_{\text{factor}}$ ) or 5% data ( $R_{\text{free}}$ ). <sup>d</sup> MolProbity reference(Williams et al., 2018)

## MATERIALS AND METHODS

### Human subjects

Blood and peripheral blood mononuclear cells (PBMCs) were isolated from COVID19+ patients using protocols approved by Institutional Review Boards at Fred Hutch Cancer Research Center, University of Washington and Seattle Children’s Research Institute.

Peripheral blood mononuclear cells (PBMCs) and serum from pre-pandemic controls were blindly selected at random from the study “Establishing Immunologic Assays for Determining HIV-1 Prevention and Control”, with no considerations made for age, or sex, participants were recruited at the Seattle Vaccine Trials Unit (Seattle, Washington, USA). Informed consent was obtained from all participants and the University of Washington and/or Fred Hutchinson Cancer Research Center and CHUM Institutional Review Boards approved the entire study and procedures.

### Recombinant Proteins

pαH-derived plasmids encoding a stabilized His- and strep-tagged SARS-CoV-2 330 ectodomain (pαH-SARS-CoV-2 S-2P), SARS-CoV-1 S-2P (pαH-SARS-CoV S-2P), MERS S2-P (pαH-MERS S-2P), SARS-CoV-2 receptor binding domain (RBD) fused to a monomeric Fc (pαH-RBD-Fc), SARS-CoV-1 RBD fused to a monomeric Fc (pαH-SARS-CoV RBD-Fc) and MERS RBD (pαH-MERS RBD-Fc) fused to a monomeric Fc have been previously described and were a kind gift from Dr. Jason McLellan (Pallesen et al., 2017; Wrapp et al., 2020b).

Proteins were produced as described in (Seydoux et al., 2020). Briefly, 1L of 293 EBNA cells at 1 x 10^6^ cells/mL were transfected with 500 mg of pαH-SARS-CoV-2 S2P, pαH-SARS-CoV S2P, pαH-SARS-CoV-2 RBD-Fc, pαH-SARS-CoV RBD-Fc, pαH-MERS S2P, pαH-MERS RBD-Fc using 2 mg of polyethyleneimine (Polyscience, Cat# 24765). After 6 days of growth, supernatants were harvested and filtered through a 0.22 mM filter. S2P supernatants were passed over a HisTrap FF affinity column (GE Healthcare, Cat# 17-5255-01) and further purified using a 2 mL StrepTactin sepharose column (IBA Lifesciences Cat# 2-1201-002) and a Strep-Tactin Purification Buffer Set (IBA Lifesciences Cat#2-1002-001). The S-2P variants were further purified using a Superose 6 10/300 GL column. RBD proteins were purified using protein A agarose resin (Goldbio CAT# P-400), followed by on-column cleavage with HRV3C protease (made in-house) to release the RBD from the Fc domain. The RBD containing flow through was further purified by SEC using a HiLoad 16/600 Superdex 200 pg column (GE healthcare). Proteins were flash frozen and stored at −80°C until use.

HCoV-OC43, HCoV-HKU1, HCoV-NL63 and HCoV-229E S1+S2 ECTs (Cat#’s 40607-V08B, 40606-V08B, 40604-V08B, 40605-V08D), SARS-HCoV-2 S1 domain (CAT#: 40591-V08B1), SARS-CoV-2 S1 N-terminal domain (CAT#40591-V41H) SARS-HCoV-2 S2 extra-cellular domain (CAT#: 40590-V08B) and SARS-CoV2 RBD (CAT#: 40150-V05H) were purchased from SinoBiologicals.

### Cell Lines

All cell lines were incubated at 37°C in the presence of 5% CO2. 293-6E (human female, RRID:CVCL_HF20) and 293T cells (human female, RRID:CVCL_0063) cells were maintained in Freestyle 293 media with gentle shaking. HEK-293T-hACE2 (human female, BEI Resources Cat# NR-52511) were maintained in DMEM containing 10% FBS, 2 mM L-glutamine, 100 U/ml penicillin, and 100 µg/ml streptomycin (cDMEM).

### ELISA

S-2P and RBD were coated onto 384-well nunclon plates (Thermo Fisher Scientific) at 0.5 mg/mL in 30 ml overnight at 4C. Plates were washed with PBS 0.02% Tween (wash buffer) using a Biotek 405 select plate washer and then blocked in 100 mL of 10% milk, 0.02% Tween (Blocking/Dilution buffer) for 1 hour at 37C. Plates were washed again, and sera was loaded at a starting dilution of 1:50 with 11 serial 1:3 dilutions in dilution buffer in a total volume of 30 mL. After another hour at 37C, plates were washed again, and IgG, IgA or IgM was detected with 30 mL of HRP secondary (Goat anti-human IgG HRP, Goat anti-human IgA HRP, Goat anti-human IgM HRP, all Southern biotech) at a 1:3000 dilution for 1 hour at 37C. After the last wash, plates were developed with 30 mL SureBlue TMB Microwell Peroxidase Substrate (Seracare KPL). The reaction was quenched with 30 mL of 1N sulfuric acid. Plates were read on a SpectraMax M2 (Molecular Devices) plate reader at 450 nM.

### B cell sorting

B cell sorting was performed as described in (Seydoux et al., 2020). Briefly, fluorescent probes were made from SARS-CoV-2 S-2P and RBD. S-2P and RBD were biotinylated protein at a theoretical 1:1 ration using the Easylink NHS-biotin kit (Thermo Fisher Scientific) according to manufacturer’s instructions. Excess biotin was removed via size exclusion chromatography using an Enrich SEC 650 10 x 300 mm column (Bio-Rad). The S-2P probes were made at a ratio of 2 moles of trimer to 1 mole streptavidin, one labeled with phycoerythrin (PE) (Invitrogen), and one with brilliant violent (BV) 711 (Biolegend), both probes were used in order to increase the specificity of detection and reduce identification of non-specific B cells. The RBD probe was prepared at a molar ratio of 4 moles of protein to 1 mole of Alexa Fluor 647-labeled streptavidin (Invitrogen). PBMCs from the five participants were thawed and stained for SARS-CoV-2-specific IgG+ memory B cells. First, cells were stained with the three SARS-CoV-2 probes for 30 minutes at 4°C, then washed, and stained with: viability dye (7AAD, Invitrogen), CD14 PE-Cy5, CD69 APC-Fire750, CD8a Alexa Fluor 700, CD3 BV510, CD27 BV605, IgM PE-Dazzle594 (BioLegend), CD4 brilliant blue 515 (BB515), IgD BV650, IgG BV786, CD56 PE-Cy5, CD19 PE-Cy7, and CD38 PerCP-Cy5.5 (BD Biosciences) for another 30 minutes at 4°C. The cells were washed twice and resuspended for sorting in 10% FBS/RPMI containing 7AAD. Cells were sorted on a FACS Aria II (BD Biosciences) by gating on singlets, lymphocytes, live, CD3-, CD14-, CD4-, CD19+, IgD-, IgG+, S-2P-PE+ and S-2P-BV711+. 10-18 million PBMCs were sorted from each participant with 384-1736 S-2P++ B cells sorted **(Supplemental Table 1)**. Cells were sorted into 96-well plates containing 16 μl lysis buffer (3.90% IGEPAL, 7.81 mM DTT, 1250 units/ml RNase Out).

### PCR amplification and sequencing of VH and VL genes

RNA was reverse transcribed to cDNA by adding 4 ul of iScript (Bio-Rad, Cat#: 1708891) to sorted B-cells and cycling according to the manufacturer’s instructions. VH and VL genes were amplified using two rounds of PCR. First round reactions contained 5 ul cDNA, 1-unit HotStarTaq Plus (QIAGEN, Cat#: 203607), 190 nM 3’ primer pool, 290 nM 5’ primer pool, 300 μM GeneAmp dNTP Blend (Thermo Fisher Scientific, Cat#: N8080261), 2 ul 10x buffer, and 12.4 ul nuclease-free H2O. Second round PCR reactions used 5 ul first round PCR as template and 190 nM of both 5’ and 3’ primers. Second round PCR products were subjected to electrophoresis on a 1.5% agarose gel containing 0.1% Gel Red Nucleic Acid Stain (Biotium, Cat#: 41002). Positive wells were then purified using either ExoSAP-IT (Affymetrix, Cat#: 78201) following manufacturer’s instructions or using a homemade enzyme mix of 0.5 units Exonuclease I (NEB, Cat#: M0293S), 0.25 units of rAPid Alkaline Phosphatase (Sigma, Cat#:4898133001), and 9.725 ul 1x PCR buffer (Qiagen) mixed with 5 ul of second round PCR product and cycled for 30 minutes at 37C followed by 5 minutes at 95C. Purified samples were Sanger sequenced (Genewiz, Seattle, WA). IMGT/V-QUEST was used to assign V, D, J gene identity, and CDRL3 length to the sequences (Brochet et al., 2008). Sequences were included in analysis if V and J gene identity could be assigned and the CDR3 was in-frame.

### VH and VL cloning and antibody production

For sorts CV1, CV2 and CN, paired VH and VL sequences were optimized for human expression using the Integrated DNA Technologies (IDT) codon optimization tool. Sequences were ordered as eBlocks (IDT) and cloned into full-length pTT3 derived IgL and IgK expression vectors (Snijder et al., 2018) or subcloned into the pT4-341 HC vector (Mouquet et al., 2010) using InFusion cloning (InFusion HD Cloning Kit, Cat#: 639649).

Sorts CV3 and PCV1 were directly cloned using Gibson Assembly. Second round PCR primers were adapted to include homology regions that corresponding to the leader sequence and constant regions on the expression vector. Cycling parameters and post-PCR clean-up remained the same. The backbone expression plasmid was amplified using primers specific for the leader sequence and constant regions in 25 μl reactions containing 2x Platinum SuperFi II DNA polymerase (Invitrogen, Cat# 12358010), 100 nM 5’ and 3’ primers, 10 ng template DNA, and 21.5 μl Nuclease-free water. The reaction was cycled at 98C for 30 seconds, 30 cycles of 98C for 10 seconds, 60C for 10 seconds, and 72C for 3 minutes and 30 seconds, followed by 72C for 5 minutes. The reaction was treated with 20 units of dpnI (NEB, Cat#:R0176S) and incubated at 37C for 60 minutes. The reaction was purified using a PCR clean-up kit according to manufacturer’s directions (NEB, Cat#: T1030S) or using ExoSAP-IT. The cloning reaction was performed using 100 ng of second round PCR product, 25 ng of backbone, 1 μl 5x InFusion HD Enzyme and nuclease-free water for a total reaction volume of 3 ul and incubated at 50C for 15 minutes.

The cloning reactions were used to transform OneShot DH5 Alpha cells (Thermo Fisher Scientific, Cat#: 12297016) according to manufacturer’s directions and plated on agar plates containing ampicillin and grown overnight. Colonies were used to seed 5 mL LB broth cultures containing ampicillin. DNA was prepared using QIAprep Spin Miniprep Kit (Qiagen, Cat#: 27106). Equal amounts of heavy and light chain expression plasmids and a 1:3 ratio of PEI was used to transfect 293-6E cells at a density of 1×10^6 cells/mL in Freestyle 293 media (Thermo Fisher Scientific, Cat#: 12338018). Supernatants were collected 6 days post transfection by centrifugation at 4,000g followed by filtration through a 0.22 μM filter (Thermo Fisher Scientific, Cat#: SE1M179M6). Clarified supernatants were then incubated with Protein A agarose beads (Thermofisher, Cat#: 20334) overnight followed by extensive washing with 1x PBS. Antibodies were eluted using 0.1M Citric Acid into a tube containing 1M Tris then buffer exchange into 1xPBS using an Amicon centrifugal filter (Thermo Fisher Scientific, Cat#: UFC901024). 198 monoclonal antibodies (mAbs) were ultimately produced and characterized. We recently reported an initial characterization of the anti-S antibody responses generated by CV1 (Seydoux et al., 2020).

### 10X sequencing

PBMCs were thawed in a 37°C water bath with pre-warmed RPMI + 10% FBS. B cells were isolated from all samples using the EasySep Human B cell Isolation Kit (Cat #17954). For the B cell receptor sequencing, cells were partitioned into gel-bead-emulsions and a cDNA was generated with each cell carrying a unique 10x identifier using the Chromium Single Cell 5’ Library and Gel Bead Kit (Cat#1000014) and the Chromium Single Cell A Chip Kit (Cat #1000009). The cDNA was enriched for V(D)J cDNA using the Human B cell Chromium Single Cell V(D)J Enrichment Kit (Cat#1000016) followed by library construction to add the priming sites used by Illumina sequencers. The V(D)J enriched library was sequenced on an Illumina HiSeq or MiSeq. Data was analyzed using the Loupe V(D)J Browser (v. 3.0.0). 15,000 cells were analyzed per donor yielding 5,000-7,000 clonotypes each. Fred Hutch Genomics core performed the sequencing and the Fred Hutch Bioinformatics core performed processing of the raw sequence data.

### BLI

All BLI experiments were performed on an Octet Red instrument at 30°C with shaking at 500-1000 rpm. All loading steps were 300s, followed by a 60s baseline in KB buffer (1X PBS, 0.01% Tween 20, 001% BSA, and 0.005% NaN_3_, pH 7.4), and then a 300s association phase and a 300s dissociation phase in KB. For the binding BLI experiments, mAbs were loaded at a concentration of 20 mg/mL in PBS onto Anti-Human IgG Fc capture (AHC) biosensors (Fortebio). After baseline, probes were dipped in either SARS-CoV2 proteins; SARS-CoV-2 RBD, S-2P, S1, S1 NTD orS2; SARS-CoV proteins; SARS-CoV-RBD or S-2P, or human coronavirus spike proteins; HCoV2-OC43, HKU1, NL63 or 229, at a concentration of 2-0.5 mM for the association phase. The binding of mature VRC01 was used as negative control to subtract the baseline binding in all of these experiments.

### ACE2 competition BLI

To measure competition between mAb and RBD for ACE2 binding, ACE2-Fc was biotinylated with EZ-Link NHS-PEG4-Biotin (Thermo Fisher Scientific) at a molar ratio of 1:2. Biotinylated protein was purified using a Zeba spin desalting column (Thermo Fisher Scientific). ACE2-Fc was then diluted to 20-83.3 mg/mL in PBS and loaded onto streptavidin biosensors (Forte Bio). Following the baseline phase, association was recorded by dipping into a 0.5 mM solution of either SARS-CoV-2 RBD or 0.5 mM solution of SARS-CoV-2 RBD plus mAb. The binding of RBD and mAb to uncoated sensors was used as background binding and was subtracted from each sample. The area under the curve (AUC) of competition was compared to the AUC of the RBD-alone condition. Samples that showed reduced binding are considered competition. Some samples appear to show enhanced binding in the presence of ACE2, perhaps because ACE2 binding stabilizes and exposes their binding sites, these antibodies are considered not competitive with ACE2.

### mAb competition BLI

To measure competition between individual mAbs for binding to SARS-CoV-2 S-2P and RBD, S-2P and RBD were biotinylated using EZ-Link NHS-PEG4 Biotin at a molar ratio of 1:2/ Biotinylated protein was purified using a Zeba spin desalting column. RBD was loaded onto streptavidin biosensors. For these experiments, following the baseline in KB, the probe was dipped in the first mAb for a first association phase, with this mAb at a saturating concentration of 2 mM. This was followed by a second baseline in KB. The probe was then dipped into the secondary mAb, at a concentration of 0.5 mM for a second association phase, followed by the standard dissociation phase. For a background control, one sample was run with the second mAb identical to the first mAb, to show the residual binding capacity, and this was subtracted from all samples.

To calculate the competition percentage, the binding of the secondary antibodies to RBD or S-2P was also assessed. Here, streptavidin probes were loaded with biotinylated S-2P or RBD, probes were then dipped in the secondary antibody at 0.5 mM for the association phase, before dissociation in KB as normal. As a background control, the binding of mature VRC01 to the RBD or S-2P was assessed and subtracted from all samples. To calculate competition percentage, the area under the curve (AUC) of this binding curve was calculated, along with the AUC of the competition curve. Percent competition was calculated as: AUC binding -AUC competition x 100. Full competition was considered when less than 15% binding capacity remained.

### Neutralization Assays

HIV-1 derived viral particles were pseudotyped with full length wild-type SARS CoV-2 S (Crawford et al., 2020; Seydoux et al., 2020). Briefly, plasmids expressing the HIV-1 Gag and pol (pHDM-Hgpm2), HIV-1Rev (pRC-CMV-rev1b), HIV-1 Tat (pHDM-tat1b), the SARS CoV2 spike (pHDM-SARS-CoV-2 Spike) and a luciferase/GFP reporter (pHAGE-CMV-Luc2-IRES-ZsGreen-W) were co-transfected into 293T cells at a 1:1:1:1.6:4.6 ratio using 293 Free transfection reagent (EMD Millipore Cat #72181) according to the manufacturer’s instructions. The culture supernatant was harvested after 72 hours at 32°C, clarified by centrifugation, filtered and frozen at −80C.

293 cells stably expressing ACE2 (HEK293T-hACE2) were seeded at a density of 4000 cells/well in a 100 µl volume in flat clear bottom, black walled, tissue culture 96-well plates. The next day, mAbs were initially diluted to 10 or 100 µg/ml in 60 µl of cDMEM in 96 well round bottom plates in duplicate, followed by a 3-fold serial dilution. An equal volume of viral supernatant was added to each well and incubated for 60 min at 37C. Meanwhile 50 µl of cDMEM containing 6 µg/ml polybrene was added to each well of 293T-ACE2 cells (2 µg/ml final concentration) and incubated for 30 min. The media was aspirated from 293T-ACE2 cells and 100 µl of the virus-antibody mixture was added. The plates were incubated at 37°C for 72 hours. The supernatant was aspirated, and cells were lysed with 100 µl of Steadyglo luciferase reagent (Promega), and luminescence was read on a Fluoroskan Ascent Fluorimeter. CV1-30 was used as a positive control and AMMO 1 (Snijder et al., 2018) was used as a negative control. Control wells containing virus, but no antibody (cells + virus) and no virus or antibody (cells only) were also included on each plate.

% neutralization for each well was calculated as the RLU of the average of the cells + virus wells, minus test wells (cells +mAb + virus) and dividing this result difference by the average RLU between virus control (cells+ virus) and average RLU between wells containing cells alone, multiplied by 100. The antibody concentration that neutralized 50% of infectivity (IC_50_), or serum dilution that neutralized 50% infectivity (ID_50_)was interpolated from the neutralization curves determined using the log(-inhibitor) versus response-variable slope (four parameters) fit using automatic outlier detection in GraphPad Prism software.

The neutralizing activities of CV1-1 and CV1-30 mAbs were also determined with a slightly different pseudovirus-based neutralization assay as previously described (Bottcher et al., 2006; Naldini et al., 1996).

### Monitoring RBD-binding to 293-ACE2 cells by flow cytometry

8 pmol of biotinylated S-2P with strep tag peptide sequence on C terminus were mixed with 10 pmol of mAb and incubated for 10 min at RT in a round-bottom tissue culture 96-well plate. 200,000 HEK293T-hACE2 cells in 50 µL of cDMEM were then added to each well and the mixture of cells + RBD or S-2P + mAb was incubated for 20 min on ice. Samples were washed once with ice-cold FACS buffer (PBS + 2% FBS + 1 mM EDTA), before staining cells with DY-549-labeled strep-tactin (1:100 dilution, IBA lifesciences, Cat #2-1565-050) or Allphycocyanin-labeled streptavidin (1:200 dilution, Agilent, Cat #PJ27S-1). Cells were washed once with FACS buffer, fixed with 10% formalin for 15 min on ice in the dark, and resuspended in 200 μl of FACS buffer to be analyzed by flow cytometry using a LSRII (BD). Control wells were included on each plate and either had no mAb, no RBD or no S-2P, or were unstained. The mean fluorescence intensity (MFI) for each sample was determined and each sample was normalized to the MFI of the no mAb control.

### Fab purification

Antigen binding fragment (Fab) was generated by incubating IgG with LysC (New England Biolabs, Cat# P8109S) at a ratio of 1 μg LysC per 10mg IgG at 37°C for 18hrs. Fab was isolated by incubating cleavage product with Protein A resin for 1hr at RT. Supernatant containing Fab was collected and further purified by SEC.

### Crystal Screening and Structure Determination

The CV2-75 Fab and SARS-CoV-2 RBD complex was obtained my mixing Fab with a 2-fold molar excess of RBD and incubated for 90 min at RT with nutation followed by SEC. The complex was verified by SDS-PAGE analysis. The complex was concentrated to 19 mg/mL for initial crystal screening by sitting-drop vapor-diffusion in the MCSG Suite (Anatrace) using a NT8 drop setter (Formulatrix). Initial crystal conditions were optimized using the Additive Screen (Hampton Research, HR2-138) Diffracting crystals were obtained in a mother liquor (ML) containing 0.1 M Tris, pH 7.5, 0.1 M Calcium Acetate, 15% (w/v) PEG 3350, and 4mM glutathione. The crystals were cryoprotected by soaking in ML supplemented with 30% (v/v) ethylene glycol. Diffraction data was collected at Advanced Photon Source (APS) SBC 19-ID at a 12.662 keV. The data set was processed using XDS (Kabsch, 2010) to a resolution of 2.80Å. The structure of the complex was solved by molecular replacement using Phaser (McCoy et al., 2007) with a search model of SARS-CoV-2 RBD (PDBid: 6xe1) (Hurlburt et al., 2020) and the Fab structure (PDBid: 4fqq) (Mouquet et al., 2012) divided into Fv and Fc portions. Remaining model building was completed using COOT (Emsley and Cowtan, 2004) and refinement was performed in Phenix (Adams et al., 2004). The data collection and refinement statistics are summarized in **Supplemental Table 3**. Structural figures were made in Pymol.

### Negative-stain EM

SARS-2 CoV 6P S protein was incubated with a three-fold molar excess of CV1-1 Fab for 30 minutes at room temperature. The complex was diluted to 0.03 mg/ml in 1X TBS pH 7.4 and negatively stained with Nano-W on 400 mesh copper grids. For data collection, a Thermo Fisher Tecnai Spirit (120 kV) and an FEI Eagle (4k x4k) CCD camera were used to produce 296 raw micrographs. Leginon (Suloway et al., 2005) was used for automated data collection and resulting micrographs were stored in Appion (Lander et al., 2009). Particles were picked with DogPicker (Voss et al., 2009) and processed in RELION 3.0 (Scheres, 2012).

### Infection of k18-hACE2 mice with SARS-CoV-2

B6.Cg-Tg(K18-ACE2)2Prlmn/J mice were purchased from Jackson Laboratories. All mice used in these experiments were females between 8 −12 weeks of age. icSARS-CoV-2 virus (Xie et al., 2020) was diluted in PBS to a working concentration of 2 x 10^5^ pfu/mL. Mice were anesthetized with isoflurane and infected intranasally with icSARS-CoV-2 (50 uL, 1 x 10^4^ pfu/ mouse) in a ABSL-3 facility. Mice were monitored daily for weight loss. All experiments adhered to the guidelines approved by the Emory University Institutional Animal Care and Committee. At the indicated day post infection, mice were euthanized via isoflurane overdose and lung tissue was collected in Omni-Bead ruptor tubes (VWR, 10032-358) filled with 1% FBS-HBSS or Tri Reagent (Zymo, #R2050-1-200). Tissue was homogenized in an Omni Bead Ruptor 24 (5.15 ms, 15 seconds). To perform plaque assays, 10-fold dilutions of viral supernatant in serum free DMEM (VWR, #45000-304) were overlaid on Vero-hACE2/TMPRSS2 monolayers and adsorbed for 1 hour at 37°C. After adsorption, 0.8% Oxoid Agarose in 2X DMEM supplemented with 10% FBS (Atlanta Biologics) and 5% sodium bicarbonate was overlaid, and cultures were incubated for 72 hours at 37°C. Plaques were visualized by removing the agarose plug, fixing the cell monolayer for 15-30 min in 4% paraformaldehyde, and staining with a crystal violet solution (20% methanol in ddH_2_O). RNA was extracted from Tri Reagent using a Direct-zol RNA MiniPrep Kit (Zymo, #R2051), then converted to cDNA using the High-capacity Reverse Transcriptase cDNA Kit (Thermo Fisher Scientific, #4368813). RNA levels were quantified using the IDT Prime Time Gene Expression Master Mix, and Taqman gene expression Primer/Probe sets (IDT). All qPCR was performed in 384-well plates and run on a QuantStudio5 qPCR system. SARS-CoV-2 viral RNA-dependent RNA polymerase levels were measured as previously described (Vanderheiden et al., 2020). The following Taqman Primer/Probe sets (Thermo Fisher Scientific) were used in this study: Gapdh (Mm99999915_g1).

### Sequence analysis

Sequences were analyzed using Geneious software (Version 8.1.9). Identification and alignments to VH/VL genes, quantification of mutations and CDRH3 length were done using V Quest (Brochet et al., 2008). Mutations were counted beginning at the 5’ end of the V-gene to the 3’ end of the 428 FW3.

### Statistical Analysis

All graphs were completed using GraphPad Prism. For column analysis of multiple independent groups one-way ANOVA with Tukey’s multiple comparison test or with Dunnett’s multiple comparison test was used. For grouped analysis two-way ANOVA with Tukey’s or Šídák’s multiple comparison test. Correlations were determined using nonparametric spearmen correlation and p values and nonlinear fit R squared values are reported.. *p<0.05, **p<0.01, ***p<0.001, ****p<0.0001.

